# Cross-family small GTPase ubiquitination by the intracellular pathogen *Legionella pneumophila*

**DOI:** 10.1101/2023.08.03.551750

**Authors:** Adriana M. Steinbach, Varun L. Bhadkamkar, David Jimenez-Morales, Erica Stevenson, Gwendolyn M. Jang, Nevan J. Krogan, Danielle L. Swaney, Shaeri Mukherjee

## Abstract

The intracellular bacterial pathogen *Legionella pneumophila* (*L.p.*) manipulates eukaryotic host ubiquitination machinery to form its replicative vacuole. While nearly 10% of *L.p.*’s arsenal of ∼330 secreted effector proteins have been biochemically characterized as ubiquitin ligases or deubiquitinases, a comprehensive measure of temporally resolved changes in the endogenous host ubiquitinome during infection has not been undertaken. To elucidate how *L.p* hijacks ubiquitin signaling within the host cell, we undertook a proteome-wide analysis of changes in protein ubiquitination during infection. We discover that *L.p.* infection results in increased ubiquitination of host proteins regulating subcellular trafficking and membrane dynamics, most notably 63 of ∼160 mammalian Ras superfamily small GTPases. We determine that these small GTPases predominantly undergo non-degradative monoubiquitination, and link ubiquitination to recruitment to the *Legionella*-containing vacuole membrane. Finally, we find that the bacterial effectors SidC/SdcA play a central, but likely indirect, role in cross-family small GTPase ubiquitination. This work highlights the extensive reconfiguration of host ubiquitin signaling by bacterial effectors during infection and establishes simultaneous ubiquitination of small GTPases across the Ras superfamily as a novel consequence of *L.p.* infection. This work positions *L.p.* as a tool to better understand how small GTPases can be regulated by ubiquitination in uninfected contexts.

## Introduction

*Legionella pneumophila* (*L.p.*) is an intracellular bacterial pathogen that has proved to be a master manipulator of its eukaryotic hosts. It is the causative agent of Legionnaires’ disease, a severe pneumonia that affects immunocompromised patients upon exposure to contaminated aerosols. In the context of human disease, *L.p.* infects alveolar macrophages, but its preferred hosts include a wide range of protozoa, demonstrating the bacterium’s ability to manipulate conserved eukaryotic processes to promote pathogenesis (Best and Kwaik, 2018; Gomez-Valero and Buchrieser, 2019). Phagocytosis by a permissive host cell triggers a complex pathogenic program in which *L.p.* avoids clearance by the endolysosomal system and instead remodels its plasma membrane-derived phagosome into an endoplasmic reticulum (ER)-like compartment called the *Legionella*-containing vacuole (LCV) (Hubber and Roy, 2010). Pathogenesis is mediated by an enormous arsenal of over 330 bacterial proteins (“effectors”) injected into the host cell cytosol by a type IV secretion system (T4SS) called Dot/Icm. Characterization of effector function has revealed numerous host targets, including membrane trafficking, autophagy, translation, and protein homeostasis (Qiu and Luo, 2017; Lockwood *et al*., 2022). Despite these advances, many aspects of *L.p.*-mediated pathogenesis remain elusive, including the functions and targets of most effectors. Studying the effects of these proteins on host cell pathways offers a great potential for the discovery of novel pathogenic and cell biological mechanisms.

Among the many host cell proteins targeted by *L.p.*, small GTPases in the Ras superfamily have long been of interest. Small GTPases are found across eukaryotes, and subfamily members regulate essential cellular functions such as cell proliferation (e.g., Ras), intracellular membrane traffic (e.g., Rab, Arf), cytoskeletal structure (e.g., Rho, Rac), and nuclear import/export (e.g. Ran) (Cherfils and Zeghouf, 2013). Despite having disparate cellular functions, these proteins share a similar bimodal activity cycle: an active, membrane associated, GTP-bound state that allows for the interaction with GTPase-specific binding partners, and an inactive, cytosolic, GDP-bound state. The small GTPase activity cycle is highly regulated – GDP release is mediated by guanine nucleotide exchange factors (GEFs), and GTPase activity and subsequent inactivation is stimulated by GTPase activating proteins (GAPs) (Cherfils and Zeghouf, 2013). GTPase activity, membrane association, and binding interactions can be further regulated by post-translational modifications (PTMs), creating an additional layer of modular control (Homma *et al*., 2021; Lei *et al*., 2021; Osaka *et al*., 2021). Given the essential roles small GTPases play in the eukaryotic cell and the diversity of regulatory mechanisms used to control GTPase function, pathogens often target GTPases through direct binding interactions and post-translational modifications (Aktories and Schmidt, 2014), and *L.p.* is no exception. The activity of small GTPases in the early secretory pathway, including Arf1, Sar1, and Rab1, has long been associated with formation of the LCV (Kagan and Roy, 2002; Derré and Isberg, 2004; Kagan *et al*., 2004). In addition, numerous effectors have been characterized with the ability to bind or post translationally modify various small GTPases, as well as recruit or remove small GTPases from the LCV membrane (Nagai *et al*., 2002; Machner and Isberg, 2006; Murata *et al*., 2006; Ingmundson *et al*., 2007; Müller *et al*., 2010; Mukherjee *et al*., 2011; Schoebel *et al*., 2011). Developing an understanding of how small GTPases are regulated during *L.p.* infection has informed a broader understanding of GTPase membrane targeting determinants as well as GTPase regulation via PTMs (Goody *et al*., 2017), positioning *L.p.* well as a tool to interrogate small GTPase regulatory mechanisms.

Another central element of *L.p.* pathogenesis is the manipulation of host cell ubiquitin signaling (Luo *et al*., 2021). Ubiquitin is a small, highly conserved, globular protein employed as a post-translational modification to regulate a multitude of eukaryotic cellular processes, including protein degradation/turnover, cell cycle, innate immune signaling, and endocytosis (Komander and Rape, 2012; Yau and Rape, 2016). Ubiquitin is covalently attached to substrate protein lysines using ATP and the sequential activity of ubiquitin-activating (E1), ubiquitin-conjugating (E2), and ubiquitin ligase (E3) enzymes, and can be removed by deubiquitinating enzymes (DUBs). Lysines can be modified with a single ubiquitin (mono-ubiquitination) or with polymeric ubiquitin chains (poly-ubiquitination), resulting in a vast array of regulatory outcomes depending on the site of ubiquitination, the ubiquitin chain length, and the linkage pattern of the ubiquitin chain that is formed (Komander and Rape, 2012).

Almost 30 translocated *L.p.* effectors have been characterized to possess either ubiquitin ligase or deubiquitinase activity – a remarkable fact considering that ubiquitin is a eukaryotic protein (Luo *et al*., 2021). These include the paralogous ligases SidC and SdcA, which promote the recruitment of as yet unknown ubiquitinated substrates and ER-membranes to the LCV (Luo and Isberg, 2004; Ragaz *et al*., 2008; Hsu *et al*., 2014). SidC/SdcA also play a role in the ubiquitination of two small GTPases important for *L.p.* pathogenesis, Rab1 and Rab10, although how SidC/SdcA are involved and the consequences of ubiquitination on Rab1/10 are not yet known (Horenkamp *et al*., 2014; Jeng *et al*., 2019; Liu *et al*., 2020). The repertoire of secreted ubiquitin ligases also includes the SidE family (SidE, SdeA, SdeB, SdeC), which catalyze non-canonical phosphoribosyl-ubiquitination, entirely bypassing the host E1-E2-E3 cascade (Bhogaraju *et al*., 2016; Qiu *et al*., 2016). A growing list of *L.p.* DUB effectors includes LotC/Lem27, which may regulate the deubiquitination and recruitment of Rab10 (Liu *et al*., 2020). The tight relationship between *L.p.* pathogenesis and ubiquitin has been further uncovered by studies that have connected host ubiquitin pathways to efficient translocation of effectors through the Dot/Icm T4SS (Ong *et al*., 2021), ubiquitin binding to the activation of the effector VpdC involved in vacuolar expansion (Li *et al*., 2022), and effector secretion to the suppression of ubiquitin-rich DALIS structures involved in antigen presentation by immune cells (Ivanov and Roy, 2009).

Thus far, one study has attempted to develop a global understanding of changes in the host ubiquitinome during infection using a proteomic approach. This study importantly revealed that *L.p.* utilizes the ubiquitin-proteasome system to downregulate innate immunity pathways and mTOR signaling during infection (Ivanov and Roy, 2013). However, the proteomic approach used relied on stable cell lines expressing tagged ubiquitin, which are prone to non-specific ubiquitination (Emmerich and Cohen, 2015; Peng *et al*., 2017). Modern ubiquitinomics approaches instead rely upon diGlycine enrichment, which can be used to detect endogenous ubiquitination events in the absence of tagged ubiquitin overexpression (Xu *et al*., 2010; Mertins *et al*., 2013). This technique has been employed to perform global analyses of host cell ubiquitinome changes during *Salmonella* Typhimurium and *Mycobacterium tuberculosis* infections (Fiskin *et al*., 2016; Budzik *et al*., 2020), but has not yet been used for *L.p.*-infected cells. In addition, because distinct subsets of effectors function during early and late stages of *L.p.* infection (Oliva *et al*., 2018), a dynamic, temporal profile of host protein ubiquitin changes has been needed to more deeply understand the regulatory mechanisms at play during infection. We set out to provide an unbiased global analysis of ubiquitin dynamics during *L.p.* infection, identifying key proteins and processes targeted during *L.p.* infection for ubiquitination and deubiquitination.

To identify proteins with changing ubiquitination status across the span of *L.p.* infection, we undertook a proteome-wide analysis of protein ubiquitination at 1– and 8-hours post-infection using diGlycine enrichment and mass spectrometry. Additionally, we quantified protein abundance for the pre-enriched samples as a quality control, and to identify potential degradative versus non-degradative signaling ubiquitination. Strikingly, we discovered that at least 63 of approximately 160 mammalian small GTPases across all subfamilies are ubiquitinated, but not degraded, during infection in a process dependent upon bacterial effector secretion. Importantly, a growing body of work has found that many small GTPases in the Ras superfamily can be regulated via ubiquitination outside of the context of infection, resulting in profound impacts on their activity (Lei *et al*., 2021). This suggests that *L.p.* may co-opt existing host regulatory mechanisms to control small GTPase function for its own benefit – an exciting prospect, given that the mechanisms and consequences of ubiquitination remain poorly defined for many small GTPases. Additionally, the degree of simultaneous cross-family small GTPase ubiquitination observed in our proteomics is, to our knowledge, unprecedented. We determine that small GTPase ubiquitination during *L.p.* infection is predominantly non-degradative monoubiquitination. Using the small GTPases Rab1, Rab5, and Rab10 as test cases, we demonstrate that robust recruitment of these GTPases to the LCV membrane is a requirement for their ubiquitination. We find that effectors SidC and SdcA promote but are not sufficient for Rab5 ubiquitination. Intriguingly, SidC/SdcA are also required for Rab5 recruitment to LCV, suggesting a complex interplay between SidC/SdcA activity, small GTPase membrane association, and ubiquitination. Finally, we find that SidC/SdcA are required for ubiquitination of small GTPases beyond the Rab subfamily, including RhoA, Arf1, and HRas. Altogether, our data suggest that *L.p.* modulates small GTPase activity during infection with prolific, cross-family small GTPase ubiquitination, and position *L.p.* as a tool to better understand how small GTPases can be regulated by ubiquitination in uninfected contexts.

## Results

### *L.p.* infection induces T4SS-dependent ubiquitinome changes in the host cell

To identify host cell components and pathways targeted with ubiquitin during *L.p.* infection, we performed a global proteomics analysis of protein ubiquitination changes in *L.p.*-infected cells. We chose HEK293 cells stably expressing the FcγRIIb receptor (HEK293 FcγR cells), as HEK293 FcγR have been used extensively in previous studies of *L.p.* pathogenesis and efficiently internalize antibody-opsonized *L.p*. (Mukherjee *et al*., 2011; Treacy-Abarca and Mukherjee, 2015; Qiu *et al*., 2016; Black *et al*., 2019; Moss *et al*., 2019). Cells were left uninfected or infected with either wild-type (WT) *L.p.* or the non-pathogenic *L.p.* Δ*dotA* strain (**Fig 1A**). For temporal resolution, infected cells were lysed at 1– or 8-hours post infection (hpi). Extracted proteins from these five conditions (uninfected control, WT 1hr, WT 8hr, Δ*dotA* 1hr, Δ*dotA* 8hr) were trypsinized and processed with diGlycine (diGly) remnant enrichment, which is found upon protein modification with ubiquitin. While diGly enrichment also captures peptides modified with the ubiquitin-like proteins NEDD8 and ISG15, these peptides make up only a small fraction of the total enriched pool (∼5%) (Kim *et al*., 2011). It is important to note that this enrichment strategy can identify only canonically ubiquitinated sites; phosphoribosyl ubiquitination mediated by the SidE family will not be detected, nor can ubiquitin chain length at a detected site be determined. Enriched peptides were then subjected to mass spectrometric analysis and quantified with appropriate adjustments made based on quality control metrics (see Materials and Methods, **Supplemental Table 1**, **Fig 1-S1A-B**). Peptide intensities between all three biological replicates per condition showed a robust reproducibility with correlation coefficients ranging from 0.80 to 0.91 (**Fig 1-S1C**). To capture the overall similarities and differences between the five experimental conditions, we performed a Principal Component Analysis (PCA). PCA identified a larger correlation between uninfected control and Δ*dotA* relative to WT (**Fig 1-S1D**). This indicates that, as expected, most changes in the ubiquitinome during infection are driven by effector secretion from *L.p.* WT.

**Figure 1:**
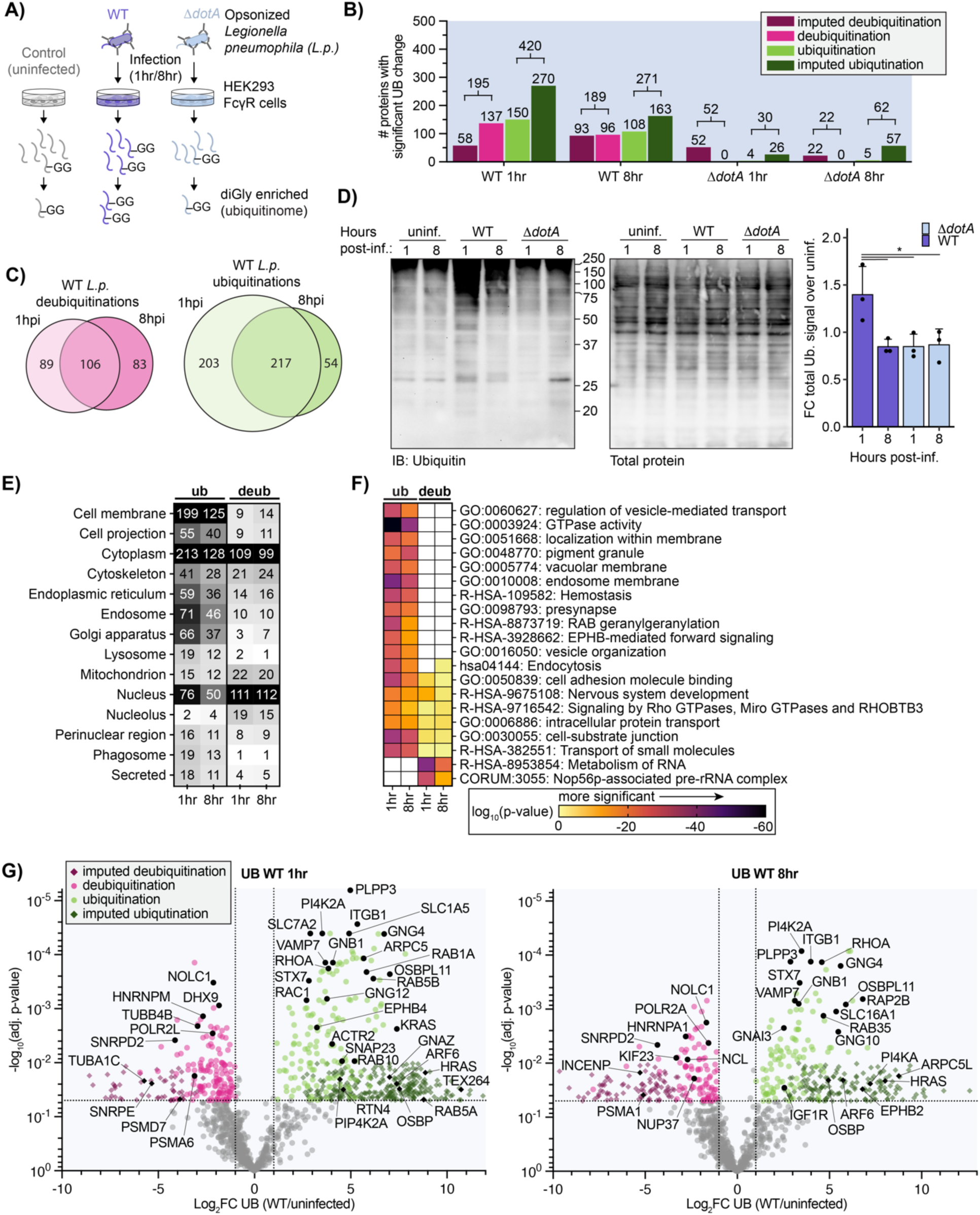
*L.p.* infection induces T4SS-dependent ubiquitinome changes in the host cell. (**A**) Schematic of experimental procedures. **(B**) Counts of proteins with a significant increase (green) or decrease (magenta) in ubiquitination compared to uninfected control for the indicated infection conditions. (Significance threshold for all subsequent analysis: |log2(FC)|>1, p<0.05, see note in text on use of imputation in this dataset). **(C**) Overlap of proteins with a significant increase (green) or decrease (magenta) in ubiquitination compared to uninfected control in the 1-hour vs 8-hour WT *L.p*. infected conditions. **(D**) Immunoblot analysis of the total pool of ubiquitinated proteins in HEK293T FcγR cells infected with WT or Δ*dotA L.p.* for 1 or 8 hours, or left uninfected. Invitrogen No-Stain protein labeling reagent was used to quantify total protein before immunoblot analysis. Total ubiquitin signal was first normalized to total protein for each sample, then the fold change over the appropriate uninfected sample was calculated. Data was subjected to a one-way ANOVA followed by a post-hoc Tukey-Kramer test for pairwise comparisons (* = p<0.05, n=3). **(E**) Subcellular localization analysis of proteins with a significant increase or decrease in ubiquitination compared to uninfected control during WT *L.p*. infection for 1 or 8 hours. **(F**) Pathway and protein complex analysis of proteins with a significant increase or decrease in ubiquitination compared to uninfected control during WT *L.p*. infection for 1 or 8 hours. Terms not significantly enriched for a given experimental condition are represented by white boxes. Analysis performed using Metascape (see methods). **(G**) Volcano plot representation of all ubiquitinome data in WT vs uninfected comparison at 1– and 8-hours post-infection. Imputed values are shown as diamonds. Significance threshold is indicated by the dotted line.

We next determined how ubiquitination was changing for individual proteins between the different conditions. We calculated the Log2 fold changes (Log2FC), corresponding p-values, and adjusted p-values for all detected proteins across all pairwise combinations of conditions (uninfected, WT and Δ*dotA* infected). Unsurprisingly, we encountered many instances in which a peptide was uniquely detected in one of the conditions while missed in the other one (e.g., a novel protein ubiquitination detected in WT infected but not uninfected control cells). Log2FC and adjusted p-values were calculated for these events using a suitable imputation strategy in which the missing peptide intensity value was assigned from the threshold of detection (see Methods). The full dataset for changes in protein ubiquitination, as well as a dataset containing changes at specific diGly sites (ubiquitin sites tab) can be found in **Supplemental Table 2.** In our subsequent analyses, we focused on four comparisons: WT1hr-Control, WT8hr-Control, Δ*dotA*1hr-Control, and Δ*dotA*8hr-Control (hereafter referred to as WT1hr, WT8hr, Δ*dotA*1hr, and Δ*dotA*8hr). Significant ubiquitination was determined using joint thresholds of |Log2FC| ≥ 1, adj.-p-value < 0.05.

Using these significance criteria, we undertook an analysis of changes in host protein ubiquitination during WT and Δ*dotA* infections. In accordance with the strong WT *L.p.*-induced ubiquitination signature shown by our PCA, we detected hundreds of proteins with significant ubiquitination changes during WT *L.p.* infection, in stark contrast to the few changes induced during Δ*dotA* infection (**Fig 1B**). The number of ubiquitinated proteins was highest early in WT *L.p.* infection, with 420 proteins ubiquitinated at 1hpi and 271 at 8hpi. In addition, we note that 80% (217 of 271) proteins ubiquitinated at 8hpi were also ubiquitinated at 1hpi, demonstrating a high degree of overlapping ubiquitination at early and late timepoints (**Fig 1C**). Analysis of total ubiquitinated proteins during infection by Western blotting confirms our proteomic data, showing significantly higher levels of ubiquitinated proteins during WT infection compared to Δ*dotA*, as well as a decrease in ubiquitination at 8hpi (**Fig 1D**).

To better understand which subcellular compartments were most targeted with ubiquitination or deubiquitination, we used subcellular localization identifiers from UniProt to tabulate the number of significantly regulated proteins per compartment (**Fig 1E**). In addition, we performed biological pathway and protein complex enrichment (**Fig 1F, Supplemental Table 3**). Subcellular localization analysis demonstrated the ubiquitination of hundreds of proteins with cytosolic or cell membrane localization, as well as endosomes, the endoplasmic reticulum, and the Golgi apparatus (**Fig 1E**). Closer study of the enrichment results reveals increased ubiquitination in pathways supporting secretory and endocytic membrane trafficking, cytoskeletal dynamics, and membrane biology (**Fig 1F**). Several of the most strongly enriched terms related to small GTPases and GTPase activity, namely “GTPase activity” and “RAB geranylgeranylation”. Further analysis of ubiquitinated proteins revealed proteins in almost all subfamilies of the Ras superfamily of small GTPases, including RAB, RAS, RHO/RAC, RAN, ARF/SAR GTPases (**Fig 1G, Supplemental Table 2**). While *L.p.* effectors are known to manipulate several of these small GTPases during infection, including Arf1, Rap1, Rab1, Rab10, Rab33b, and Ran (Nagai *et al*., 2002; Rothmeier *et al*., 2013; Qiu *et al*., 2016; Schmölders *et al*., 2017; Steiner *et al*., 2018; Jeng *et al*., 2019; Liu *et al*., 2020), the targeting of numerous small GTPases with ubiquitination is unprecedented. Regulatory small GTPase ubiquitination is known to occur in uninfected contexts (Lei *et al*., 2021), suggesting possible widespread manipulation of small GTPase signaling during *L.p.* infection. GTPase ubiquitination during WT infection extended to numerous heterotrimeric G proteins (**Fig 1G**). This included alpha (GNA11/13/I1/I2/I3/O1/Q/Z), beta (GNB1/2/4), and gamma (GNG4/5/7/10/12) subunits, as well as several regulators of heterotrimeric G protein signaling (RGS17/19/20). Although the role of heterotrimeric G protein signaling has not been studied extensively in the context of *L.p.* infection, it is known that G proteins are important for phagocytosis of *L.p.* in amoeba (Fajardo *et al*., 2004) and that multiple G proteins are found on the surface of the LCV in proteomic datasets (Hoffmann *et al*., 2014). Ubiquitination of heterotrimeric G protein subunits can result in a wide variety of signaling outcomes (Dewhurst *et al*., 2015; Torres, 2016; Dohlman and Campbell, 2019), suggesting *L.p.* or the host cell may modify G protein signaling via ubiquitination during infection.

We also detected ubiquitination on other proteins or pathways known to be targeted by *L.p.* effectors but not known to be targeted with ubiquitination. These include numerous regulators of the actin cytoskeleton (ARPC1B/2/5/5L, ACTR2/3, BAIAP2), proteins involved in lipid exchange (OSBP, OSBPL3/8/9/11, PITPNA), lipid kinases (PI4KA, PI4K2A, PIP4K2A), as well as SNARES and membrane fusion regulators (STX3/6/7/10/12, SNAP23/29, VAPA, NAPA, VAMP7). Also ubiquitinated during infection were several proteins known to be modified with non-canonical ubiquitination by the SidE family of *L.p.* effectors – which is not detected by the diGly enrichment technique used here – including the ER-shaping proteins RTN4, FAM134C, and TEX264 (Shin *et al*., 2020). In addition, we identified ubiquitination on protein targets previously unknown to play roles in *L.p.* infection, including solute carrier transporters, tyrosine (EPHB1/2/4, FGFR2, IGF1R) and serine/threonine-protein kinases (LIMK1, PKN2, TNIK, MINK1), and integrins (ITGA3/B1/B1BP1/B3).

In addition to protein ubiquitination, *L.p.* infection induced the deubiquitination of hundreds of proteins at both 1hpi (195 proteins) and 8hpi (189 proteins) (**Fig 1B**). Of these, 106 proteins were deubiquitinated at both timepoints, suggesting that early and late infection deubiquitination is targeted to many of the same proteins (**Fig 1C**). Unlike the strong ubiquitination of cell membrane proteins, proteins deubiquitinated during WT infection primarily localized to the nucleus and the cytoplasm (**Fig 1E**). Pathway enrichment analysis of deubiquitinated proteins showed minimal overlap with pathways targeted by ubiquitination, suggesting that protein populations targeted for ubiquitination and deubiquitination during infection are distinct (**Fig 1F**). In line with a distinct, nuclear-enriched deubiquitination response, enrichment analysis primarily described deubiquitinated proteins with the two terms “Metabolism of RNA” and “Nop56p-associated pre-rRNA complex”. These enrichments are driven in part by deubiquitination of numerous spliceosome proteins (HNRNPA1/C/K/M/U, SNRPD2/D3/E, SF3B3/B6), as well as transcription regulators (DHX9, POLR2A/2L) and the multifunctional proteins nucleolin (NCL) and NOLC1 (**Fig 1G**). Although none of these proteins are known targets of *L.p.* effectors, host cell transcription is known to be modulated by the *L.p.* effectors LegAS4/RomA (Rolando *et al*., 2013), LphD (Schator *et al*., 2023), LegA3/AnkH (Dwingelo *et al*., 2019), and SnpL (Schuelein *et al*., 2018) through a variety of mechanisms, suggesting that *L.p.* may employ additional effectors to target nuclear function. Intriguingly, we also observed the deubiquitination of numerous subunits of the proteasome (PSMA1/A6/B3/C1/C3/C5/D1/D7, ADRM1), which is known to be important for *L.p.* infection (Dorer *et al*., 2006; Price *et al*., 2011). We also noticed deubiquitination of several regulators of the RAN GTPase (RANBP2, RCC1) and associated proteins such as nuclear pore complex (NUP37/85/188, TPR), tubulin subunits (TUBA1C, TUBB, TUBB4B), the microtubule stabilizer CKAP2, and microtubule associated proteins (PCM1, INCENP, KIF23, CEP131). The deubiquitination of these proteins is intriguing because *L.p.* is known to activate the Ran GTPase – promoting microtubule polymerization and LCV motility – with the effectors LegG1 and PpgA (Rothmeier *et al*., 2013; Simon *et al*., 2014; Swart *et al*., 2020). Altogether, *L.p.* induces the deubiquitination of hundreds of proteins over the course of infection on a population distinct from proteins targeted with ubiquitination.

In contrast to WT infection, Δ*dotA* induced few changes in both protein ubiquitination and deubiquitination at both 1 and 8hpi (**Fig 1B, 1-S2**). The few changes that did occur during Δ*dotA* infection primarily occurred in the nucleus, cytoplasm, and cell membrane (**Fig 1-S2D**), and were described by terms known to relate to bacterial infection such as “Bacterial invasion of epithelial cells”, “PID NFkappaB Canonical Pathway”, “lytic vacuole” and “PCP/CE pathway” (planar cell polarity pathway) (**Fig 1-S2E**) (Tran *et al*., 2014). The enrichment of these terms during Δ*dotA* infection indicates a strong antibacterial host response which is absent during WT *L.p*. infection, and serves as a confirmation that our proteomic analysis aligns with the biology of the system.

Given the tight relationship between protein ubiquitination and degradation, we compared host cell protein ubiquitin changes to changes in abundance. To do this, we analyzed our pre-diGly enriched cell lysates via mass spectrometry and quantitated changes in host protein abundance. As with our ubiquitinomics, peptide intensities showed robust reproducibility and PCA distinctly separated WT infected cells from uninfected and Δ*dotA* infected cells (**Fig 1-S3**). Log2 fold changes, corresponding p-values, and adjusted p-values for all detected proteins across all pairwise combinations of conditions were computed and analyzed and can be found in **Supplemental Table 2 (abundance tab**) and **Fig 1-S4A-D.** To compare protein ubiquitination and abundance changes, we plotted ubiquitination Log2FC values against abundance Log2FC for all detected proteins at 1hpi and 8hpi (**Fig 1-S4E**). We used the same significance cutoff of |Log2FC| ≥ 1, adj.-p-value < 0.05 to determine proteins significantly changing in abundance, ubiquitination, or both abundance and ubiquitination. Importantly, few proteins experienced both significant abundance and ubiquitination changes simultaneously. This result serves as a quality control that changes in abundance are not responsible for detected changes in ubiquitination, and suggests that ubiquitination largely does not result in protein abundance changes during infection.

### *L.p.* infection results in the ubiquitination of multiple Ras superfamily small GTPases

Among the many ubiquitin-regulated pathways and proteins during infection, we were particularly intrigued by the ubiquitination of many small GTPases in the RAS superfamily. Previous studies have shown that *L.p.* uses ubiquitin to modify select small GTPases in the Rab subfamily. Two Rab proteins known to play important roles in pathogenesis, Rab1 and Rab10, are ubiquitinated during infection in a process dependent upon the paralogous effectors SidC and SdcA (Kagan *et al*., 2004; Horenkamp *et al*., 2014; Jeng *et al*., 2019). In addition, Rab33b is targeted for non-canonical phosphoribosyl ubiquitination by the SidE family of ligases (Qiu *et al*., 2016). Although the consequences of Rab1/10/33b ubiquitination are not known, both SidC/SdcA and SidE family effectors are associated with timely LCV formation, suggesting that *L.p.* may ubiquitinate small GTPases for pathogenic benefit. Additionally, although small GTPases are known to be regulated by ubiquitination outside the context of infection, the simultaneous cross-family ubiquitination of these proteins is unprecedented and suggests that *L.p.* may exploit a GTPase regulatory mechanism common to the entire superfamily (Lei *et al*., 2021). Thus, we further investigated *L.p.*-induced cross-family small GTPase ubiquitination to learn more about *L.p.* pathogenesis, but also GTPase regulation more broadly.

We first identified the number and family range of small GTPases ubiquitinated during infection in our ubiquitin site dataset (**Supplemental Table 2 – ubiquitin_sites tab**). Small GTPases in the Ras superfamily accounted for 132 of 868 significant ubiquitination sites (15.21%) at 1hpi, and 77 of 532 (14.47%) significant ubiquitination sites at 8hpi (**Fig 2A**, **Fig 2-S1**). Ubiquitination sites were detected on at least 63 of the approximately 163 known mammalian Ras superfamily small GTPases, falling on members of the ARF, RAN, RHO/RAC, RAS, and RAB subfamilies (**Fig 2-S2**). Surprisingly, many of these ubiquitination sites were imputed, suggesting that the ubiquitination of these proteins may be catalyzed by *L.p.* effectors (**Fig 2-S2**). While several of the small GTPases ubiquitinated during infection are known to be regulated by ubiquitin outside of the context of infection, these ubiquitination events are often transient and hard to detect (Lachance *et al*., 2013; Shin *et al*., 2017; Sapmaz *et al*., 2019; Duncan *et al*., 2022), suggesting that *L.p.* infection may ubiquitinate small GTPases at a higher frequency or with a greater stability than observed in uninfected cells. In contrast to WT infection, Δ*dotA* infection induced ubiquitination of few small GTPases, consistent with mass small GTPase ubiquitination being an effector-induced process.

**Figure 2:**
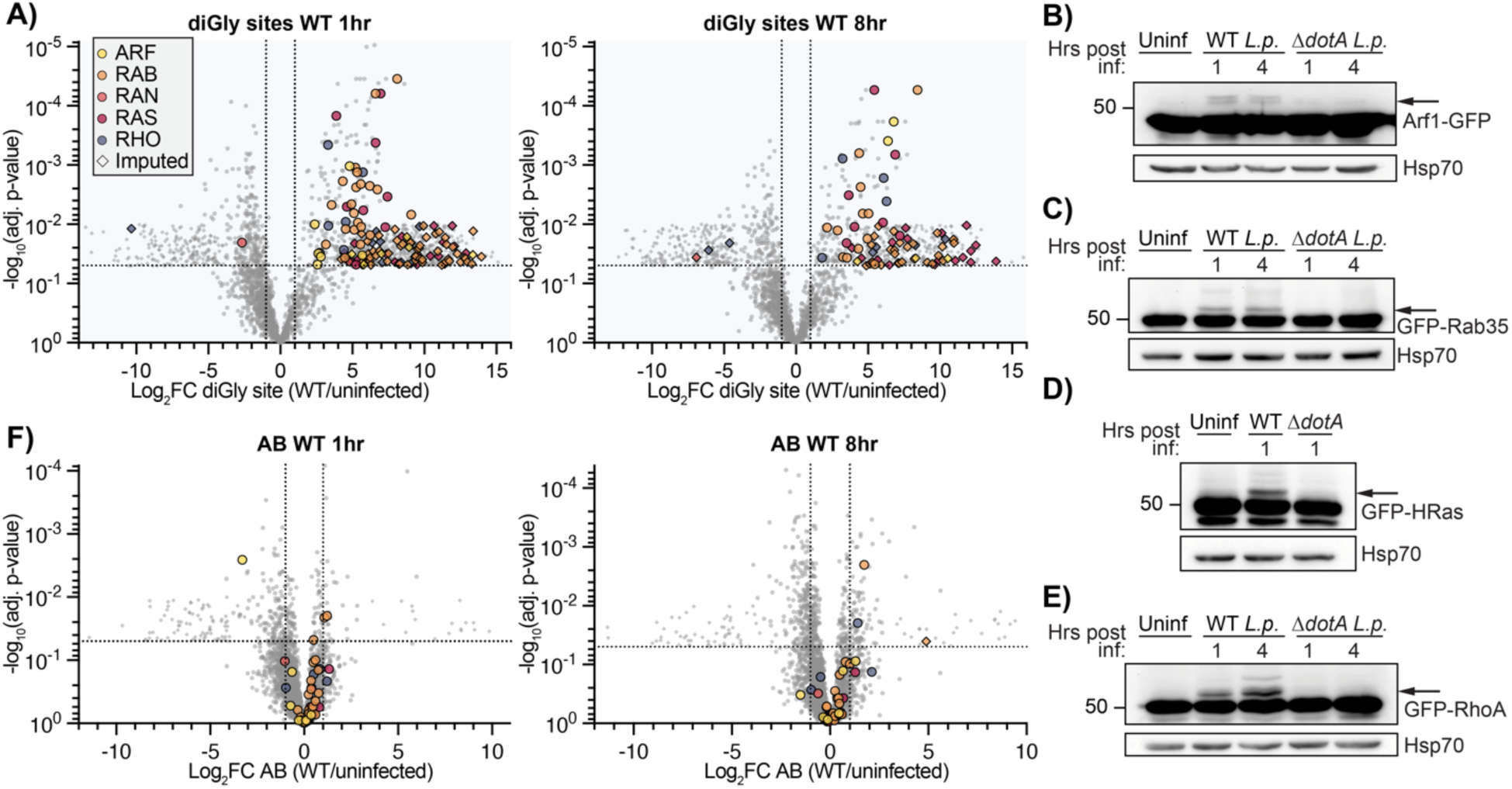
Small GTPases across the Ras superfamily are mono-ubiquitinated during WT *L.p.* infection. (**A**) Volcano plot representation of diGlycine site dataset for WT *L.p.* vs. uninfected comparison at 1– and 8-hours post-infection. Each point represents a unique diGlycine enriched peptide; for some GTPases, multiple peptides were detected. Imputed values are shown as diamonds, and Ras superfamily subfamilies are differentiated by color. Significance threshold is shown by the dotted line. **(B**)-**(E**) Immunoblot analysis of lysates prepared from HEK293T FcγR cells transiently transfected with the indicated GFP-tagged small GTPase, then infected with either WT or Δ*dotA L.p.* for 1 or 4 hours, or left uninfected. Blots were probed with anti-GFP and anti-Hsp70 antibodies. Monoubiquitinated GTPase indicated with an arrow. **(F**) Volcano plot representation of abundance dataset as in (A).

Since our diGly enrichment ubiquitinomics strategy precludes determination of ubiquitin chain length, we assessed the ubiquitination of multiple Ras superfamily small GTPases via Western blot. We transfected HEK293T FcγR cells with a panel of GFP-tagged or Flag-tagged GTPases and infected them with WT or Δ*dotA L.p.*. We expected ubiquitinated GTPases to show the appearance of bands in multiples of ∼8.5kDa (molecular weight of a ubiquitin moiety) above the major, non-ubiquitinated species. Indeed, numerous GTPases in the ARF (ARF1, ARF6), RAS (HRas, Rap1, Rap2B), RHO (RhoA, RhoB, RhoC, RhoQ), and RAB (Rab6A, Rab9A, Rab20, Rab35) subfamilies showed a mass shift during WT but not Δ*dotA* infection (**Fig 2B-E**, **Fig 2-S3**), both confirming the cross-family GTPase ubiquitination observed in our mass spectrometric data and suggesting that the majority of small GTPase ubiquitination occurs as monoubiquitination. In some circumstances, we noticed multiple mass shifts above the unmodified band, as seen for RAS and RHO GTPases (**Fig 2D-E**, **Fig 2-S3C-D**). These higher molecular weight bands are consistent with the conjugation of either polyubiquitin chains or multiple monoubiquitin moieties to distinct lysine residues on these small GTPases. Regardless, these higher molecular weight bands are clearly lower abundance than the +8.5kDa shifted band, suggesting that GTPases are primarily monoubiquitinated during infection.

To determine if cross-family small GTPase ubiquitination may promote degradation, we mined our AB dataset for changes in small GTPase abundance during infection (**Fig 2F**). Of the many detected GTPases, almost all fell below both the adj. p-value and the Log2FC significance cutoffs, suggesting that GTPases do not significantly change in abundance during infection. This result is consistent with past work demonstrating that *L.p.*-induced Rab1 ubiquitination is removed at later time points during infection in a proteasome-independent process (Horenkamp *et al*., 2014), and with past work on non-degradative small GTPase monoubiquitination (Sapmaz *et al*., 2019; Kholmanskikh *et al*., 2022). Consistent with this insight from our proteomics analysis, we do not see a decrease in small GTPase abundance across the time course of infection by Western blot for all small GTPases tested (**Fig 2B-E**, **Fig 2-S3, Fig 5**).

We next decided to explore small GTPase sequence and structure for clues regarding the impacts of ubiquitination. Towards this end, we aligned the sequences of the significantly ubiquitinated GTPases, annotated regions of interest, and marked all unique ubiquitination or deubiquitination sites from both 1 and 8hpi (**Fig 3S1 – full alignment, Fig 3A – Rab1A only**). Regions of interest include: (1) the five conserved G boxes important for contact with GTP/GDP, (2) the Switch I and Switch II regions important for interaction between active GTPases and their downstream binding partners, (3) the C-terminal hypervariable domain (HVD) typically responsible for proper membrane targeting and subcellular localization, and (4) the five alpha helices and six beta sheets characteristic of most Ras superfamily small GTPases. We next defined 10 regions based on these conserved structural and functional elements. Within each region, we counted the number of WT *L.p.*-induced ubiquitin sites (**Fig 3B**). Surprisingly, most of the ubiquitination did not occur in Switch I / II regions (regions #3 and #5), known to be targeted with PTMs by many pathogens, including *L.p.*, to block interaction between active GTPases and their downstream binding partners (Aktories and Schmidt, 2014). Instead, 92 of 138 unique ubiquitination sites (∼67%) were detected within the three C-terminal regions: the α4 helix (region #8, 35 sites), G5 box lysine (region #9, 21 sites), and α5/C-terminal hypervariable domain (region #10, 36 sites) (**Fig 3B**). Intriguingly, most work on small GTPase ubiquitination in uninfected contexts has determined ubiquitination to primarily fall within these regions (Steklov *et al*., 2018; Osaka *et al*., 2021; Kholmanskikh *et al*., 2022), suggesting that *L.p.* infection hijacks GTPase regulatory regions targeted in the absence of infection as well.

**Figure 3:**
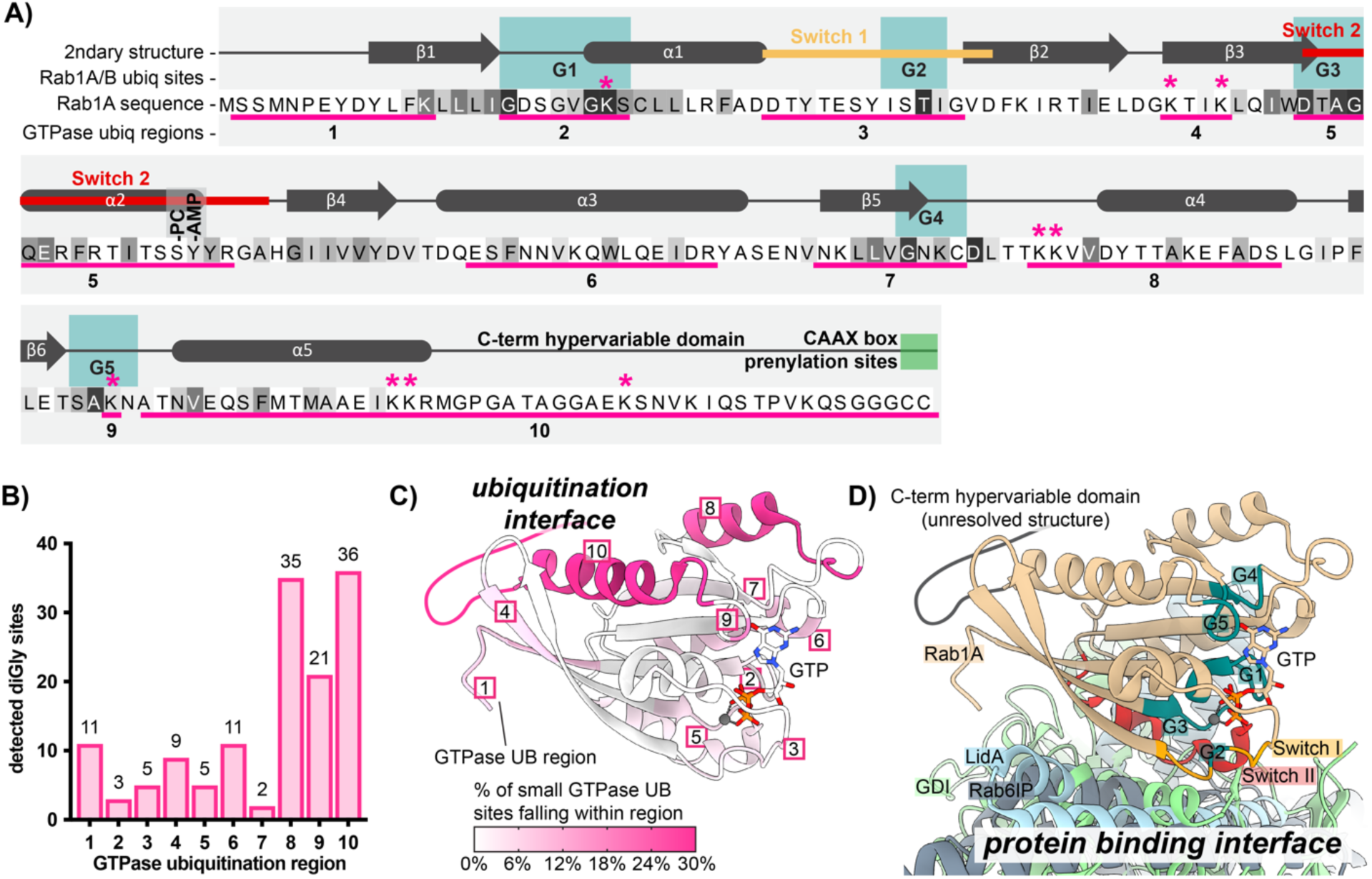
Small GTPase ubiquitinations cluster in the C-terminal region. (**A**) Schematic of small GTPase structural and functional regions, using Rab1A as an example. Regions frequently ubiquitinated across detected small GTPases are underlined in pink and numbered 1-10. Sequence colored by conservation within the small GTPase superfamily, from white (non-conserved) to black (extremely highly conserved residue); see full alignment in **Fig 3S1**. The “*” symbol indicates ubiquitination sites detected for Rab1A or Rab1B. The site of phosphocholination by *L.p*. effector AnkX (S79) is annotated as “-PC”, while the site of AMPylation by *L.p*. effector DrrA (Y80) is marked as “-AMP”. **(B**) Pooled counts of significant ubiquitinations detected in ubiquitination regions defined in (A) across all 63 ubiquitinated small GTPases. **(C**) Structure of Rab1A with the 10 structural/functional regions indicated. Regions are colored by percentage of detected small GTPase ubiquitin sites falling within a given region. **(D**) Alignment of structures for human Rab1A (tan) in complex with *L.p.* effector LidA (light blue), mouse Rab6 bound to its effector Rab6IP (gray), or yeast Rab1 homologue YPT1 bound to GDI (light green). Important structural and functional domains of Rab1A are colored and labeled. PDB accession numbers: 3TKL (Rab1A:LidA), 2BCG (YPT1:GDI), and 3CWZ (Rab6:R6IP1).

We next mapped ubiquitinated regions onto the structure of the small GTPase Rab1 (**Fig 3C, pink regions**). To visualize the relationship between these ubiquitinated regions, key functional regions, and protein binding interfaces, we also aligned the structure of Rab1 to structures of GTPases bound to several types of partners, including Rab1 bound to the *L.p.* secreted Rab-binding effector LidA, yeast YPT1 (Rab1 homologue) bound to GDI, and mouse Rab6 bound to Rab6-interacting protein 1 (R6IP1) (**Fig 3D**). As expected, LidA, GDI, and R6IP1 predominantly form contacts with GTPases around the Switch I / II regions. To our surprise, the dominantly ubiquitinated regions #8, #9, and #10, localize to the distal face of Rab1, opposite protein binding regions. This result implies that cross-family small GTPase ubiquitination may not directly block GTPase-protein binding interactions, and instead, affect other intrinsic GTPase properties, such as membrane association, GTP/GDP binding, or GTP hydrolysis.

### LCV-localized pools of Rab1 are targeted for ubiquitination

We next decided to interrogate the driving forces behind small GTPase ubiquitination during infection more directly. Based on previous studies, we hypothesized that small GTPase ubiquitination may be spatially restricted to LCV-membrane localized pools of these proteins. First, past work on Rab1 ubiquitination has shown its ubiquitination at 1hpi and deubiquitination by 8hpi, which correlates with Rab1 LCV recruitment and removal (Kagan *et al*., 2004; Ingmundson *et al*., 2007; Horenkamp *et al*., 2014). Second, the effectors SidC/SdcA are known to control both Rab10 LCV recruitment and its ubiquitination, suggesting a functional link between these two processes (Jeng *et al*., 2019; Liu *et al*., 2020). Finally, it is well established that the LCV accumulates ubiquitinated proteins throughout the first 6 to 8 hours of infection, indicating that the LCV membrane may be a site of ubiquitin ligase activity (Dorer *et al*., 2006; Ivanov and Roy, 2009).

In order to test our hypothesis, we manipulated the recruitment of Rab1 to the LCV during infection and assessed changes in Rab1 ubiquitination. Recruitment of Rab1 was manipulated by altering the activity of the *L.p.* effector DrrA (also known as SidM), which recruits Rab1 to the LCV at early timepoints during infection via the activity of a Rab1-specific GEF domain and retains Rab1 in the LCV membrane via the activity of its AMP-transferase, or AMPylation domain (Murata *et al*., 2006; Müller *et al*., 2010; Hardiman and Roy, 2014). Past work has demonstrated that a DrrA genomic deletion *L.p.* strain *ΔdrrA* displays considerably reduced Rab1 recruitment to the LCV, and that DrrA AMPylation activity is required, as complementation with AMPylation-dead DrrA D110,112A fails to rescue Rab1 recruitment (Hardiman and Roy, 2014). Consistent with Rab1 LCV recruitment and retention being tied to its ubiquitination, we found considerably reduced levels of Rab1 ubiquitination during infection with *L.p.* Δ*drrA* and *L.p.* Δ*drrA + pDrrA D110,112A* compared to *L.p.* WT (**Fig 4A-B**). Ubiquitination was rescued by complementation of *L.p.* Δ*drrA* with a plasmid expressing WT DrrA. In contrast, DrrA knockout and AMPylation mutant strains had no effect on the ubiquitination of Rab10, suggesting that DrrA does not control the ubiquitination of GTPases not targeted by its GEF domain (**Fig 4-S1A-B**).

**Figure 4:**
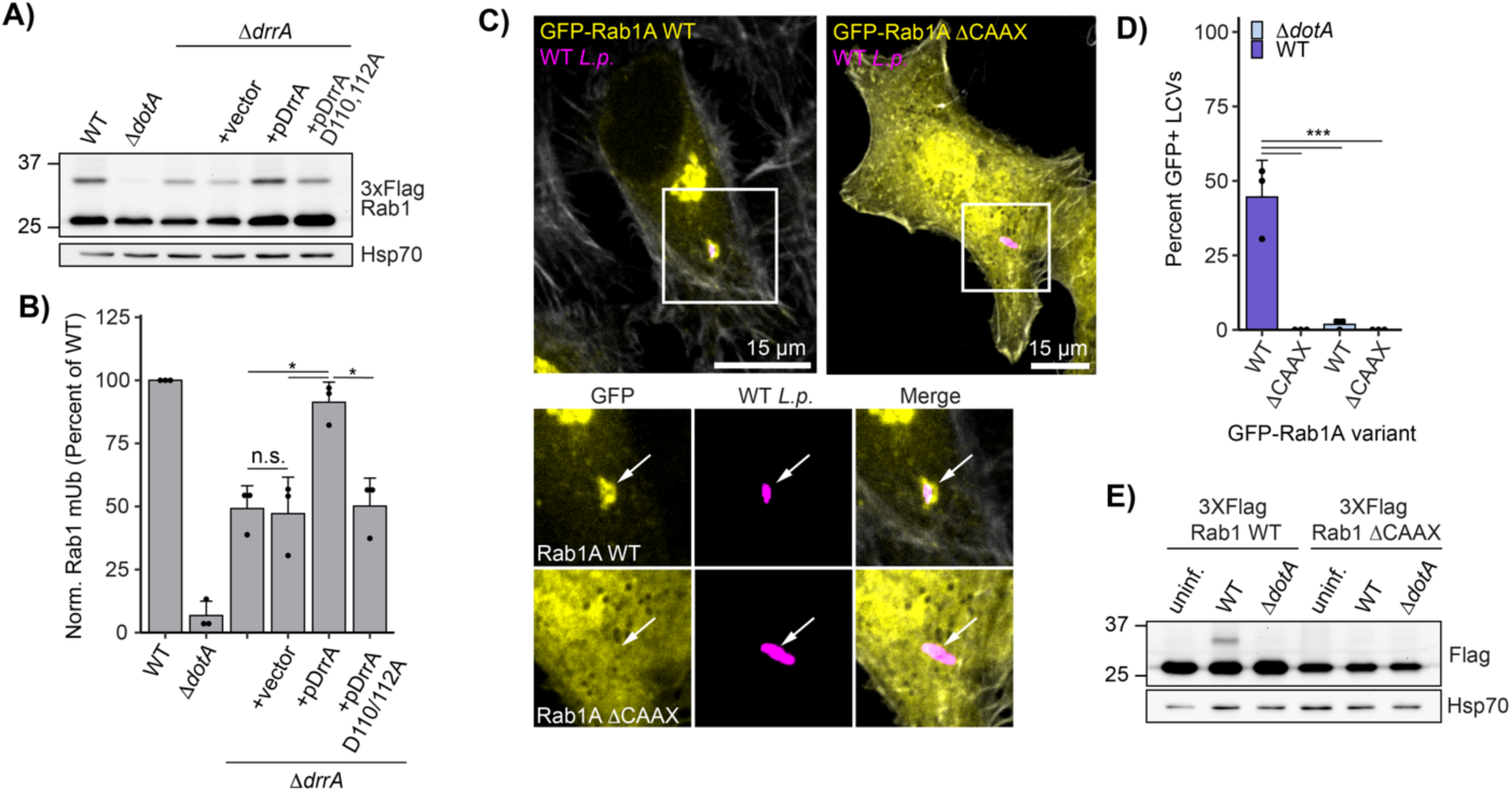
Rab1A is monoubiquitinated at the LCV membrane. (**A**) Immunoblot analysis of lysates prepared from HEK293T FcγR transfected with 3XFlag Rab1A and infected with a Δ*drrA L.p*. strain panel (WT, Δ*dotA*, Δ*drrA*, and Δ*drrA* complemented with empty vector or plasmid encoded DrrA WT or D110, 112A) for 1 hour (MOI=50). Monoubiquitinated Rab1 indicated with an arrow. **(B**) Quantification of biological replicates (N=3) of experiment shown in (A). Normalized Rab1A monoubiquitination intensity was calculated as a percentage of WT *L.p.* infection levels (see Methods). **(C)-(D**) Immunofluorescence analysis of EGFP Rab1A WT or ΔCAAX LCV recruitment. HeLa FcγR cells were transfected with indicated construct, then infected for 1 hour with either WT or Δ*dotA L.p.* (MOI=1), fixed, and stained with anti-*Legionella* antibody. **(C**) Representative images, and **(D**) quantification of EGFP positive LCVs (percent of total scored per biological replicate, n=3, 25 LCVs scored/replicate). **(E**) Immunoblot analysis of monoubiquitination of Rab1A WT vs ΔCAAX during *L.p.* infection. HEK293T FcγR cells were transfected with either 3X Flag Rab1A WT or ΔCAAX, then infected with WT or Δ*dotA L.p.* for 1 hour (MOI=50), or left uninfected. Lysates were probed with anti-Flag antibody. Monoubiquitinated Rab1 indicated with an arrow. *For all graphs:* bars represent mean value, error bars represent standard deviation. Individual points are values from each biological replicate. *Statistical analysis of Western blot quantification*: one-way ANOVA followed by Tukey-Kramer post-hoc test for each pair of means. * = p<0.05, n.s. = p>0.05.

To further interrogate the relationship between Rab1 recruitment to the LCV membrane and its ubiquitination, we sought to prevent Rab1 association with membranes entirely. To this end, we generated a lipid anchor mutant of Rab1 by deleting its two prenylation sites: the C-terminal cysteines C204 and C205 (known as the CAAX box). As expected, Rab1 ΔCAAX showed fully cytosolic localization compared to the predominantly Golgi-localized WT Rab1 and was not recruited to the LCV membrane (**Fig 4C-D**). Consistent with LCV-membrane recruitment being a prerequisite for Rab1 ubiquitination, Rab1 ΔCAAX ubiquitination was entirely abolished during infection (**Fig 4E**). We note a similar loss of ubiquitination upon deletion of the Rab10 CAAX motif (**Fig 4-S1C**). Collectively, our data suggest that only LCV-localized pools of Rab1 are targeted for ubiquitination, and that membrane recruitment may be a prerequisite for infection-induced small GTPase ubiquitination.

### Early endosomal GTPase Rab5 is recruited to the LCV and targeted for ubiquitination

We next turned to another small GTPase generally thought to be an antagonist in the *L.p.* infection cycle, Rab5 (Anand *et al*., 2020). The three genetically encoded Rab5 isoforms (Rab5A, B, and C), particularly Rab5A/B (Chen *et al*., 2009, 2014), are considered master regulators of the early endosomal compartment, recruiting proteins that direct recently endocytosed cargo within the cell and control endosomal fusion or fission (Langemeyer *et al*., 2018). Rab5 is required for endosome maturation, a remodeling of the protein and lipid components of the endosomal membrane that marks the transition from early to late endosome. Unlike early endosomes, late endosomes can fuse with lysosomes, resulting in the degradation of enclosed cargo (Langemeyer *et al*., 2018). It is well established that WT *L.p.* evades lysosomal fusion, whereas Δ*dotA L.p.* succumbs to lysosomal degradation, suggesting that bacterial effectors prevent endosome maturation at the LCV membrane (Roy *et al*., 1998; Clemens *et al*., 2000b). As such, Rab5 has been a protein of interest in the study of *L.p.* pathogenesis for years. Several bacterial effectors have been proposed to be activated by Rab5 binding (Gaspar and Machner, 2014) or regulate Rab5 activity (Sohn *et al*., 2015), but to our knowledge, none have been shown to directly post-translationally modify Rab5.

Increased ubiquitination is detected for all three Rab5 isoforms in our ubiquitinome dataset during WT, but not Δ*dotA*, *L.p.* infection (**Fig 2-S1, Supplemental Table 2**). The accumulation of a higher molecular weight species (∼8.5 kDa) is observed for endogenous Rab5A in U937 macrophage lysates across the first 6 hours of WT *L.p.* infection (**Fig 5A**). Endogenous Rab5A and Flag-tagged Rab5B and C also show this mass shift in WT *L.p.* infected HEK293T FcγR (**Fig 5-S1A**). To confirm that this higher molecular weight species is monoubiquitinated Rab5A, we performed a ubiquitin pulldown on lysates from HEK293T FcγR cells expressing Flag Rab5A and infected with *L.p.* WT or Δ*dotA*. In WT infected, but not Δ*dotA* infected or uninfected pulldown samples, we observe Flag Rab5A laddering, with a dominant band at ∼35 kDa corresponding to monoubiquitinated Rab5A (**Fig 5B**).

**Figure 5:**
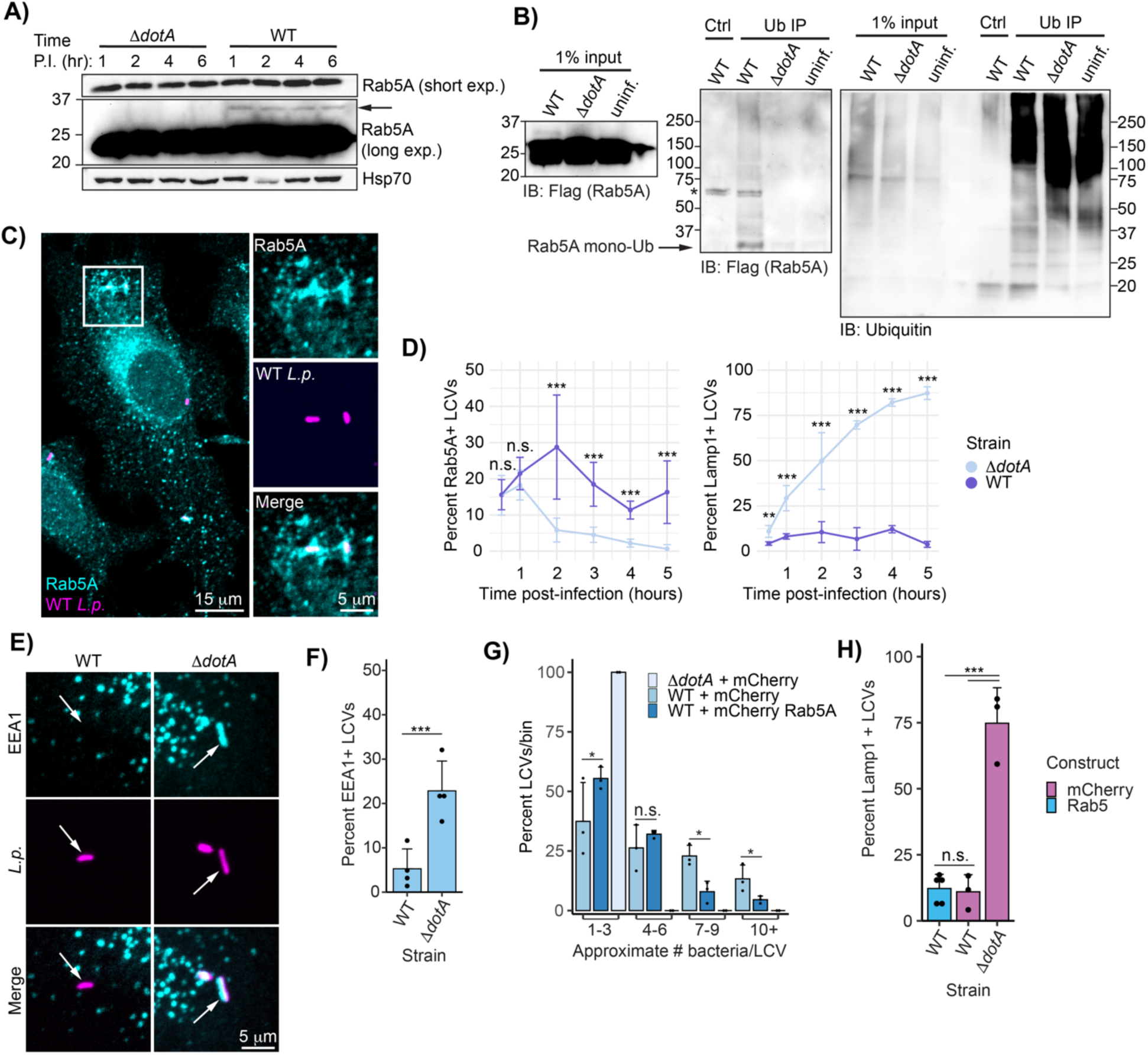
Rab5A is recruited to the LCV and ubiquitinated during WT *L.p.* infection. (**A**) Immunoblot analysis of Rab5A mass shift during WT or Δ*dotA L.p.* infection. U937 cells differentiated into macrophage-like cells were infected with either WT or Δ*dotA* and lysed at the indicated time point. **(B**) Immunoblot analysis of ubiquitin immunoprecipitation from *L.p.*-infected cells. HEK293T FcγR cells transfected with 3XFlag Rab5A were infected with WT or Δ*dotA* for 4 hours or left uninfected. Ubiquitinated proteins were enriched from these samples using ubiquitin affinity beads (SignalSeeker Kit, Cytoskeleton Inc). Input and IP samples were probed with anti-Flag and anti-ubiquitin antibodies. Asterisk (*) indicates a non-specific band. **(C**)-**(D**) Immunofluorescence analysis of Rab5A and Lamp1 LCV recruitment. HeLa FcγR cells were infected with WT or Δ*dotA* for indicated length of time, fixed, and probed with either anti-Rab5A or anti-Lamp1 antibody. **(C**) Representative image of WT LCV Rab5A recruitment (1-hour post-infection), **(D**) quantification of all experiments performed. For each biological replicate, the percent of LCVs scored positive for the indicated marker was calculated (n=3, 75-150 LCVs scored per replicate). Line graphs show the mean percent positive +/-standard deviation. **E**) Rab5 binding partner EEA1 associates with the Δ*dotA* but not WT LCV. HeLa FcγR cells were infected with *L.p.* WT or Δ*dotA* for 1 hour, fixed, and probed with anti-EEA1 and anti-*Legionella* antibodies. **F**) Quantification of biological replicates performed as in (A). For each biological replicate, the percent of LCVs scored positive for the indicated marker was calculated (n=4, 75-150 LCVs scored per replicate). **G**) *L.p.* replication is inhibited by Rab5A overexpression. HeLa FcγR cells were transfected with mCherry tagged Rab5A or mCherry alone, infected with WT or Δ*dotA* for 10 hours, fixed, and probed with anti-*Legionella* antibodies. Bacteria count per LCV was approximated by measuring the LCV area and dividing by the average area of the Δ*dotA* LCVs. For each biological replicate (n=3, 25-50 LCVs per replicate), the number of LCVs falling into the indicated bin was tabulated, and the percent each bin represented of the total was calculated. **H**) Rab5 overexpression does not increase WT LCV Lamp1 staining. HeLa FcγR cells were transfected with mCherry tagged Rab5A or mCherry alone, infected with WT or Δ*dotA* for 4 hours, fixed, and probed with anti-Lamp1 and anti-*Legionella* antibodies. For each biological replicate, the percent of LCVs scored positive for the indicated marker was calculated (n=3-4, 75-150 LCVs scored per replicate). *For all bar graphs:* bars represent mean value, error bars represent standard deviation. Individual points are values from each biological replicate. *Statistical analysis of LCV scoring quantification and replication:* G test of independence was performed on pooled counts (positive vs. negative) from all biological replicates. For multi-condition experiments, upon verifying significance (p<0.05), pairwise comparisons between strains were evaluated by post-hoc G-test using the Bonferroni-adjusted p-value as a significance threshold. P-values used: (D). 008, (F) 0.05, (G) 0.013, (H) 0.017.

Given the clear connection between ubiquitination and LCV recruitment observed for Rab1 (**Fig 4**), we next asked if Rab5A is present at the LCV at any point during infection. Previous studies disagree on whether Rab5A is recruited to the LCV during WT *L.p.* infection. Initial EM immunogold staining suggested that WT *L.p.* excludes Rab5 from the LCV membrane (Clemens *et al*., 2000a), but more recent mass spectrometry analysis of purified LCVs and immunofluorescence analysis in RAW macrophages identified Rab5 as LCV localized (Hoffmann *et al*., 2014). To address this discrepancy in the literature, we carried out immunofluorescence analysis of endogenous Rab5A in HeLa FcγR cells infected with WT or Δ*dotA L.p.* across a time range from 30 minutes to 5hpi. We observed clear Rab5A recruitment to both the WT and Δ*dotA* LCV, while the WT LCV still resists lysosomal fusion, as shown by the exclusion of the lysosomal membrane protein Lamp1 (**Fig 5C-D**). Interestingly, whereas the Δ*dotA* LCV shows more canonical Rab5A dynamics in which recruitment peaks shortly after internalization and decays quickly thereafter, the WT LCV shows moderate frequencies of Rab5A localization across the first five hours of infection. This mirrors the persistent ubiquitination observed across early infection timepoints (**Fig 5A**), suggesting that, as for Rab1, Rab5A ubiquitination requires LCV localization. To directly test this hypothesis, we generated mCherry tagged Rab5A WT and lipid anchor deletion (Rab5A ΔCAAX) constructs, and quantified localization to the WT and Δ*dotA* LCVs during infection in HeLa FcγR. Rab5A WT localizes to both the WT and Δ*dotA* LCV, whereas Rab5A ΔCAAX is diffuse and excluded from the LCV (**Fig 5-S1D-E**). Flag tagged versions of these constructs show a clear loss of monoubiquitination for the ΔCAAX construct (**Fig 5-S1F-G**), consistent with the model that Rab5A monoubiquitination requires membrane association.

We next assessed whether localization of Rab5 to the LCV results in the accumulation of early endosomal markers. Membrane-associated Rab5A recruits early endosome-specific proteins both through direct binding interactions and by the production of the phosphoinositide PI(3)P via the activity of multiple binding partners (Shin *et al*., 2005). One such protein is early endosome antigen 1 (EEA1), which binds to both active Rab5 and PI(3)P (Simonsen *et al*., 1998). Despite Rab5 association with the WT LCV, we do not observe recruitment of EEA1 at 1hpi, while EEA1 robustly localizes to the Δ*dotA* LCV (**Fig 5E-F**). Notably, multiple *L.p.* effectors are known to coordinate the conversion of PI(3)P to PI(4)P at the LCV membrane during early infection (Weber *et al*., 2006; Dong *et al*., 2016), and the exclusion of EEA1 suggests that this lipid conversion program is active even while Rab5A is present. Conversely, association of Rab5A with the WT LCV is not sufficient to promote an early endosome-like character at the membrane, in contrast to the Δ*dotA* LCV. To determine if Rab5A activity is detrimental to *L.p.*, we assayed both bacterial replication and lysosomal trafficking of the LCV in the context of Rab5 overexpression. HeLa FcγR transfected with mCherry-Rab5A or mCherry alone were infected with *L.p.* WT or Δ*dotA,* fixed at 4 and 10hpi, and subjected to immunofluorescence analysis. At 10hpi, there is a small but significant decrease in the frequency of high bacterial burden LCVs during Rab5A overexpression compared to control (**Fig 5G**). However, there is no increase in Lamp1 positive WT LCVs at 4hpi during Rab5A overexpression (**Fig 5H**), further suggesting that Rab5 recruitment and activity does not promote trafficking of the WT LCV through the endolysosomal pathway.

### Bacterial effectors SidC/SdcA are necessary but not sufficient for Rab5A monoubiquitination, and control Rab5A recruitment to the LCV

Next, we sought to identify the bacterial effectors required for Rab5A ubiquitination. Previous studies have shown that bacterial effector paralogs SidC and SdcA are required for Rab1 (Horenkamp *et al*., 2014) and Rab10 (Jeng *et al*., 2019) ubiquitination. To determine if SidC/SdcA play similar roles in Rab5 ubiquitination, we infected HEK293T FcγR cells expressing Flag Rab5A with SidC/SdcA knockout and complemented strains. Indeed, infection with a SidC/SdcA genomic deletion strain (*L.p.* Δ*sidC/sdcA*) fails to induce Rab5A monoubiquitination (**Fig 6A-B**). Transformation of the Δ*sidC/sdcA* strain with a plasmid encoding either SdcA or SidC is sufficient to rescue Rab5A monoubiquitination, suggesting that these effectors are functionally redundant in this context. SdcA/SidC have been identified as E3 ligases with unique protein folds (Hsu *et al*., 2014). While these proteins catalyze autoubiquitination *in vitro*, neither direct *in vitro* ubiquitination assays with SidC and Rab1 nor several MS based approaches have revealed host target proteins of SidC/SdcA (Hsu *et al*., 2014; Shin *et al*., 2020). In accordance with these findings, ectopic expression of SdcA/SidC in the absence of infection is not sufficient to induce Rab5A monoubiquitination (**Fig 6C**). This result suggests that the context of infection provides the complete enzymatic machinery required for SidC/SdcA-mediated Rab5 monoubiquitination, which could include either bacterial or host components, or both.

**Fig 6:**
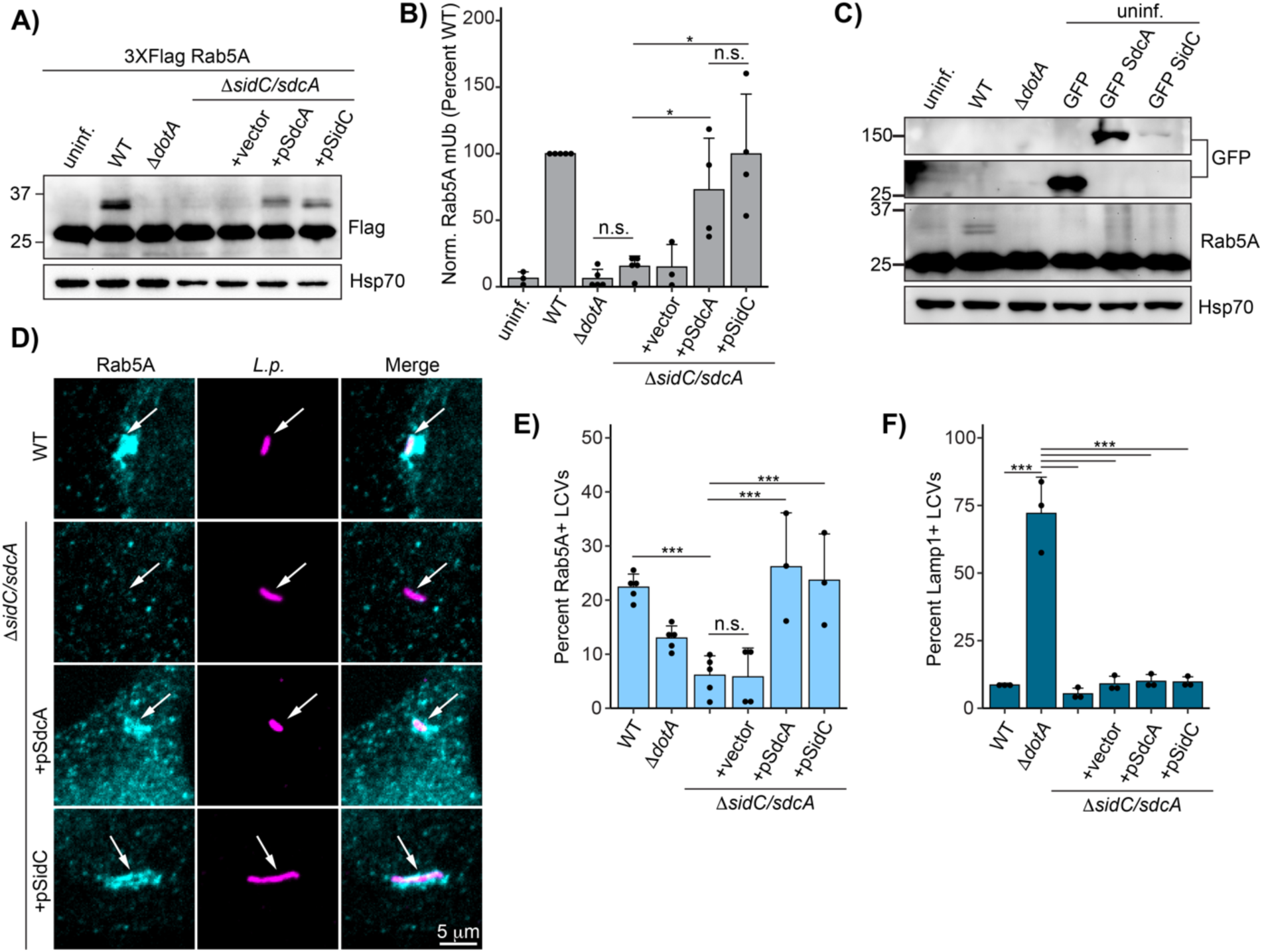
Bacterial effectors SidC/SdcA are required for Rab5A monoubiquitination and recruitment to the LCV. (**A**) Immunoblot analysis of Rab5A monoubiquitination during infection with Δ*sidC/sdcA L.p.* strain panel (WT, Δ*dotA*, Δ*sidC/sdcA*, and Δ*sidC/sdcA* transformed with vector or plasmid expressing SdcA or SidC). HEK293T FcγR cells transfected with 3XFlag Rab5A were infected with the indicated strain or left uninfected. Cells were lysed at 4 hours post infection and probed with anti-Flag antibody. **(B**) Quantification of biological replicates (N=3-5) of experiment shown in (A). Normalized Rab5A monoubiquitination intensity was calculated as a percentage of WT *L.p.* infection levels (see Methods). Data was subjected to a one way ANOVA followed by a post-hoc Tukey-Kramer test for pairwise comparisons (* = p<0.05, n=3-4). **(C**) Immunoblot analysis of Rab5A monoubiquitination during SdcA or SidC ectopic expression. HEK293T FcγR were either left untransfected (lanes 1-3) or transfected with GFP alone or GFP-tagged SdcA or SidC for 24 hours. The untransfected cells were either left uninfected or infected with WT or Δ*dotA L.p.* for 4 hours. All cells were lysed and probed with anti-Flag and anti-GFP antibodies. **(D**) Representative images of Rab5A LCV recruitment levels for the Δ*sidC/sdcA* strain panel as observed by immunofluorescence. HeLa FcγR cells were infected with indicated strain for 1 hour, fixed, and probed with anti-*Legionella* and anti-Rab5A antibodies. **(E**) Quantification of biological replicates (N=3-5) of experiment shown in (D). 75-150 LCVs were scored per replicate as positive or negative for Rab5A recruitment, and the percent Rab5A+ LCVs was calculated per replicate. **(F**) Quantification of Lamp1 LCV recruitment for the Δ*sidC/sdcA* strain panel. HeLa FcγR cells were infected with indicated strain for 4 hr, fixed, and probed with anti-*Legionella* and anti-Lamp1 antibodies. LCVs were scored as in (E). *For all graphs:* bars represent mean value, error bars represent standard deviation. Individual points are values from each biological replicate. *Statistical analysis of LCV scoring quantification:* G test of independence was performed on pooled counts (positive vs. negative) from all biological replicates. Upon verifying significance (p<0.05), pairwise comparisons between strains were evaluated by post-hoc G-test using the Bonferroni-adjusted p-value as a significance threshold (p = 0.003). * = p<0.003, ** = p<0.0003, *** = p<0.00003, n.s. = p>0.003.

As we have established a link between Rab monoubiquitination and LCV localization, we next examined whether SidC/SdcA control Rab5A recruitment. Immunofluorescence analysis reveals that the Δ*sidC/sdcA* LCV fails to accumulate endogenous Rab5A at one hour post-infection, whereas Δ*sidC/sdcA* strains complemented with either SidC– or SdcA– expressing plasmids robustly recruit Rab5A (**Fig 6D-E**). The finding that bacterial effectors recruit Rab5A is somewhat surprising, as Rab5 activity is generally thought to be deleterious to *L.p.* infection (Anand *et al*., 2020; Kim and Isberg, 2023). The Δ*sidC/sdcA* strains are as resistant to lysosomal fusion as WT *L.p.* (**Fig 6F**), consistent with a model in which Rab5 recruitment and ubiquitination are not a primary mechanism of endosome maturation subversion at the LCV membrane.

### SidC/SdcA promote small GTPase ubiquitination beyond the Rab subfamily

With the finding that SidC/SdcA regulate Rab5 ubiquitination, in addition to past work demonstrating their role in Rab1/10 ubiquitination, we hypothesized that SidC/SdcA may play a role in small GTPase ubiquitination beyond the Rab subfamily. To evaluate the role SidC/SdcA may play in cross-family small GTPase ubiquitination more broadly, we transfected HEK293T FcγR cells with GFP-tagged HRas and RhoA constructs and infected with SidC/SdcA knockout and complemented strains. Strikingly, infection with *L.p.* Δ*sidC/sdcA* abolished ubiquitination of both GTPases. Surprisingly, complementation of *L.p.* Δ*sidC/sdcA* with a plasmid encoding SidC but not SdcA rescued ubiquitination of these GTPases (**Fig 7A**). This result implicates SidC/SdcA in cross-family small GTPase ubiquitination more broadly. It also suggests that SidC/SdcA may play overlapping but distinct roles in small GTPase ubiquitination, as the ubiquitination of Rab1 seems to be primarily dependent upon the activity of SdcA (Horenkamp *et al*., 2014) (**Fig 7-S1A-B**), and the ubiquitination of Rab5 appears to be equally dependent upon SidC and SdcA (**Fig 6A-B**).

**Figure 7:**
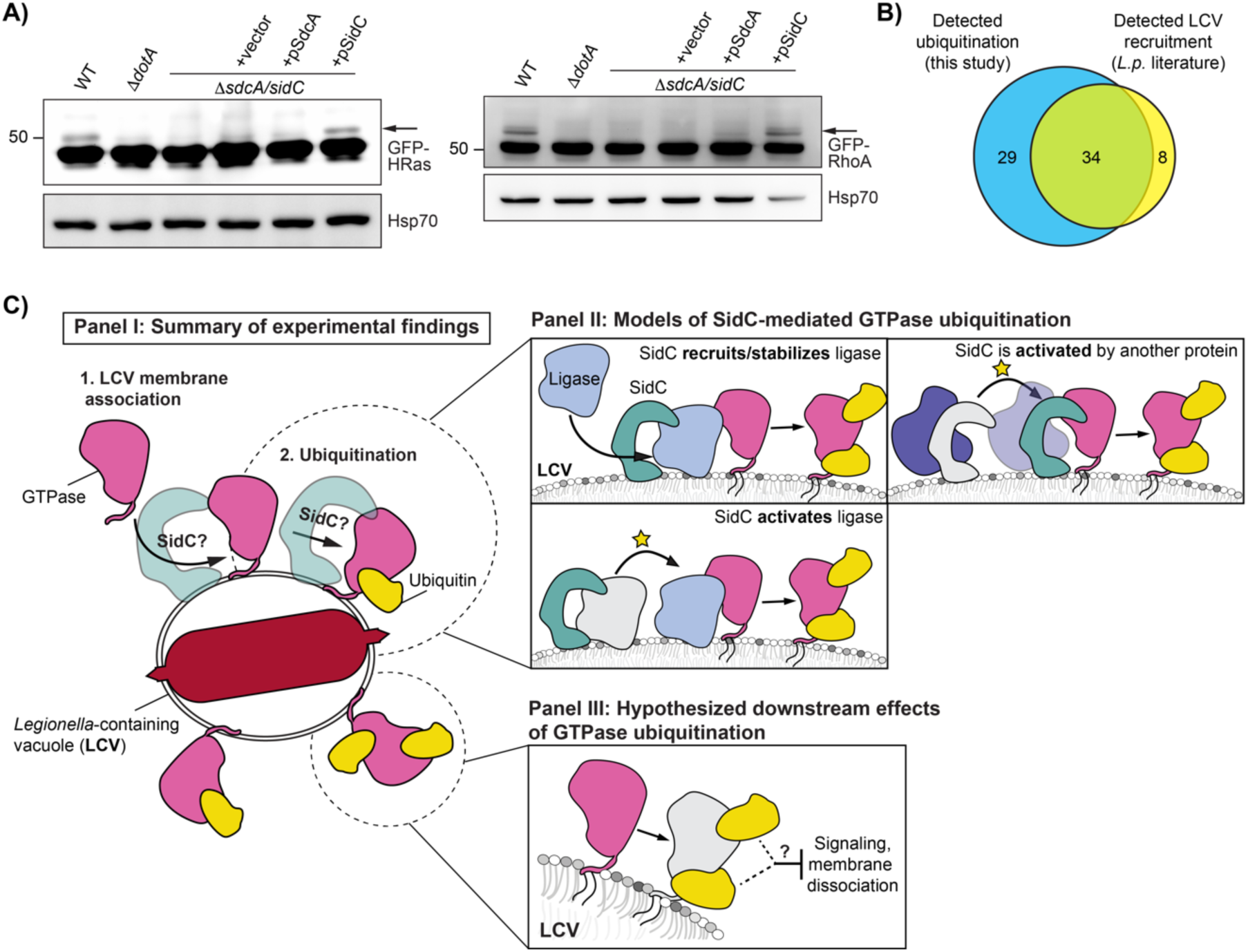
SidC/SdcA is required for small GTPase ubiquitination beyond the Rab subfamily. (**A**) Immunoblot analysis of small GTPase ubiquitination during infection with Δ*sidC/sdcA L.p.* strain panel (WT, Δ*dotA*, Δ*sidC/sdcA*, and Δ*sidC/sdcA* transformed with vector or plasmid expressing SdcA or SidC). HEK293T FcγR cells transfected with the indicated GFP-tagged small GTPase were infected (MOI=50) with the indicated strain or left uninfected. Cells were lysed at 1 hour post infection and probed with anti-GFP and anti-Hsp70 antibodies. **(B**) Comparative analysis of literature reports of small GTPase recruitment to the LCV with small GTPases found to be ubiquitinated during infection in our proteomic analysis. **(C**) Schematic of models for downstream consequences and mechanism of SidC/SdcA mediated small GTPase ubiquitination.

Our data connecting Rab1 and Rab5 ubiquitination to their LCV recruitment also strongly suggest that small GTPases that localize to the LCV are targeted for ubiquitination. To assess whether LCV localization is linked to the mass ubiquitination of small GTPases, we mined all past work on *L.p.* to generate a comprehensive list of small GTPases detected at the LCV in both mammalian and amoebal cell lines, either by immunofluorescence or purified LCV proteomic approaches (**Fig 7-S2, Fig 7-S3**). Using this approach, we determined 41 mammalian small GTPases with documented LCV localization. We find that 34 of these 42 LCV-localized GTPases overlap with the 63 ubiquitinated small GTPases in our dataset (**Fig 7B**). While by no means an exhaustive approach, this result suggests that LCV-membrane localized small GTPases are targeted for ubiquitination.

## Discussion

Here, we define the ubiquitinated proteome of HEK293 cells infected with *Legionella pneumophila* at 1– and 8-hours post infection. Analysis of this dataset reveals that infection with WT *L.p.* induces hundreds of significant changes in the host ubiquitinome spanning processes known to be involved in infection, such as membrane trafficking and lipid exchange, as well as processes with less characterized or unknown roles in infection, such as mRNA splicing and solute transport. The temporal resolution of our data highlights that the most dramatic changes in the host ubiquitinome occur at early timepoints during infection, although substantial modification of the ubiquitinome persists at 8hpi. Additionally, we see that many of the same pathways and proteins are targeted throughout infection, suggesting that similar E3 ligases and DUBs may be active throughout infection, or that many early changes in the ubiquitinome are stable. Given the connection between ubiquitination and protein degradation, we also paired our analysis of the host ubiquitinome with an analysis of changes in host protein abundance. Intriguingly, changes in ubiquitination seem to be largely independent of changes in abundance, suggesting that many of the ubiquitination changes we detected during infection are not connected to degradative signaling outcomes.

A major effect of infection was the ubiquitination of 63 of approximately 163 known small GTPases spanning RAB, RAS, RHO/RAC, RAN, and ARF/SAR subfamilies. We determined that GTPases are predominantly monoubiquitinated during infection, with the monoubiquitinated GTPase making up a small fraction of the total GTPase pool in the cell. Along with our proteomic data showing no significant small GTPase abundance changes during infection, as well as past work demonstrating that ubiquitinated Rab1 is not degraded in the proteasome (Horenkamp *et al*., 2014), these results strongly suggest that small GTPase monoubiquitination plays a non-degradative role during infection. The cross-family ubiquitination of small GTPases also appears to be specific to *L.p.* infection, as human cells infected with *Salmonella* Typhimurium or *Mycobacterium tuberculosis* do not show a comparable level of cross-family ubiquitination (Fiskin *et al*., 2016; Budzik *et al*., 2020).

Despite our determination that GTPases are primarily monoubiquitinated, our proteomic analysis identified multiple ubiquitinated lysines on most ubiquitinated GTPases. This is not entirely surprising, as ubiquitination of a given lysine residue is thought to be governed more by accessibility than amino acid sequence context; E3 ligases are often promiscuous in modifying lysine residues on substrates (Danielsen *et al*., 2011; Kim *et al*., 2011; Mattiroli and Sixma, 2014). Through sequence alignment and binning of ubiquitinated residues into different structural regions, we were able to determine that most ubiquitination sites fell within GTPase C-terminal regions after the G4 box, including the conserved G5 box SA**K** motif lysine that makes contacts with the guanine of GTP, and the hypervariable C-terminal domain (HVD), which contains sequence elements required for lipidation (Müller and Goody, 2017). Mapping these regions onto the Rab1A structure demonstrated that they form a distinct interface opposite the canonical small GTPase protein binding regions, Switch I and II. This suggests that GTPase ubiquitination during infection functions through an alternative mechanism of action compared to known PTMs within the Switch regions such as phosphorylation and AMPylation, which are known to block GTPase-protein binding interactions more directly (Müller *et al*., 2010; Tan *et al*., 2011; Aktories and Schmidt, 2014; Levin *et al*., 2016; Steger *et al*., 2016). Although several studies have investigated small GTPase ubiquitination within these regions outside the context of infection, the data on downstream consequences are mixed and appear to be highly GTPase and/or residue dependent. Monoubiquitination of RhoC, Rab11a, and KRas on either the G5 SAK motif or the preceding α4 helix appears to be activating (Sasaki *et al*., 2011; Baker *et al*., 2013; Lachance *et al*., 2013; Kholmanskikh *et al*., 2022), while ubiquitination of Rab5 in the same region appears to impair activity (Shin *et al*., 2017). Equally paradoxical, ubiquitination of Rab7 in the HVD appears to maintain it in the membrane (Sapmaz *et al*., 2019), while ubiquitination of H/N/KRas in this region prevents membrane association (Steklov *et al*., 2018). It is worth noting that GTPases can form protein-protein binding contacts outside of the Switch regions. For example, ubiquitination in the Rab7 HVD disrupts its interaction with the Rab-interacting lysosomal protein (RILP) (Sapmaz *et al*., 2019), which forms a binding interface with V180, L182, and Y183 in this region (Wu *et al*., 2005). Additionally, GTPases are also able to form key homo– and heterodimerization contacts via sites in the α3, α4, and α5 helices, which can affect their signaling (Mima, 2021). Thus, we note that the ubiquitination interface determined in our work may still directly affect small GTPase protein binding interactions.

By manipulating the recruitment of the small GTPase Rab1 to the *Legionella*-containing vacuole, we were able to determine that robust recruitment and retention of Rab1 on the LCV promotes its ubiquitination. This is most clearly demonstrated by our experiments in which we infected cells with either *L.p.* Δ*drrA* or the AMPylation mutant *L.p.* Δ*drrA* + pDrrA D110,112A. Infection with these strains substantially decreased the fraction of Rab1 monoubiquitination relative to WT *L.p.* infection, but had no impact on the monoubiquitination of Rab10, whose LCV recruitment is not controlled by DrrA. This result indicates that Rab1 recruitment is required for its ubiquitination, and that ubiquitination of other small GTPases is not contingent upon DrrA activity or Rab1 LCV association.

Paired with our Rab1 LCV-recruitment model, the observation that all Rab5 isoforms are ubiquitinated during infection led us to the finding that Rab5A is recruited to the WT LCV during infection. Previously published results conflicted on whether Rab5 associates with the WT LCV (Clemens *et al*., 2000a; Hoffmann *et al*., 2014). In the present study we relied on immunofluorescence analysis of endogenous Rab5 during infection, and found that the WT LCV stains positive for Rab5A at moderate frequencies throughout early infection. Additionally, we link Rab5 ubiquitination to LCV recruitment, and observe ubiquitination of endogenous Rab5 in U937 macrophage-like cells, suggesting that Rab5 recruitment and ubiquitination is not specific to HeLa FcγR. Surprisingly, Rab5A association with the WT LCV does not appear to be passive, as it requires the bacterial effectors SidC/SdcA, which are also necessary for Rab5A ubiquitination during infection. However, the SidC/SdcA knockout strain is as resistant to lysosomal fusion as the WT strain, suggesting that neither recruitment nor ubiquitination of Rab5 play a dominant role in evading LCV-lysosome fusion. Notably, previous reports suggest that overexpression of Rab5 antagonizes *L.p.* pathogenesis but does so by decreasing the integrity of the LCV membrane (Anand *et al*., 2020; Kim and Isberg, 2023), rather than by increasing trafficking of the LCV to the lysosome. Consistent with this finding, we observe that Rab5A overexpression results in a bacterial replication defect without an increase in Lamp1 recruitment to the WT LCV. Taken together, these results are inconsistent with a model by which Rab5 activity simply increases trafficking of the LCV to the lysosome, and instead suggest a nuanced interplay between *L.p.* effectors and Rab5 activity during infection.

Our data place the bacterial effectors SidC and SdcA at the center of cross-family small GTPase ubiquitination, although the specific role that they play is still unclear. In our experimentation, we find that SidC/SdcA are required for both Rab5A monoubiquitination and Rab5 LCV recruitment during infection. Past work has found that SidC/SdcA are also required for recruitment of Rab10 to the LCV (Jeng *et al*., 2019), but given that SidC/SdcA drive ER membrane association with the LCV (Horenkamp *et al*., 2014; Hsu *et al*., 2014), and Rab10 is ER-associated (English and Voeltz, 2013), the possibility remained that these effectors played an indirect role in bringing Rab10 to the LCV. The observation that SidC/SdcA promote the recruitment of the endosomal GTPase Rab5 in addition to the ER GTPase Rab10 suggests that these bacterial effectors may play a more direct role in cross-family small GTPase recruitment, retention, or both at the LCV membrane. Indeed, SidC/SdcA have already been linked to the recruitment of the GTPase Arf1 to the vacuole (Horenkamp *et al*., 2014), and here, we find that SidC/SdcA are required for Arf1 monoubiquitination during infection. Ubiquitination of the GTPases HRas and RhoA comparably is dependent upon the activity of SidC/SdcA. Intriguingly, we note that SidC and SdcA seem to contribute differently towards GTPase ubiquitination dependent upon the GTPase. We find that SdcA is primarily responsible for Rab1 ubiquitination, SidC is primarily responsible for Arf1, HRas, and RhoA ubiquitination, and both SidC and SdcA seem to play equivalent roles in promoting Rab5 ubiquitination. This difference in specificity implies that SidC and SdcA may target different membranes or GTPases for recruitment to the LCV, consistent with past work that has found lower conservation in the domain of SidC/SdcA hypothesized to be involved in membrane tethering.

Altogether, our data support a model in which all small GTPases in the LCV membrane are targeted with non-degradative ubiquitination on an interface distal to canonical protein binding regions. This cross-family small GTPase ubiquitination is promoted through an unknown mechanism by the effectors SidC/SdcA (**Fig 7C – Panel I**). In addition to the experiments presented here, our model is supported by the observed correlation between GTPases that have been detected on the vacuole in past literature to small GTPases with detected ubiquitination in our proteomics.

Although SidC/SdcA are known to possess ubiquitin ligase activity, our findings and past data are not consistent with a model in which SidC/SdcA directly ubiquitinate small GTPases, and instead support a model in which SidC/SdcA either recruit or activate the ligases responsible for small GTPase ubiquitination (**Fig 7C – Panel II**). Several lines of evidence support the indirect involvement of SidC/SdcA in small GTPase ubiquitination. First, ectopic expression of SidC/SdcA does not induce ubiquitination of Rab1 (Horenkamp *et al*., 2014; Hsu *et al*., 2014) or Rab5 (this study). Second, in vitro ubiquitination reactions containing purified SidC have not resulted in Rab1A ubiquitination (Hsu *et al*., 2014). Lastly, protein-protein interaction experiments have failed to detect interaction between SidC/SdcA and Rab1, Arf1, or numerous other proteins involved in LCV formation (Horenkamp *et al*., 2014). We cannot rule out the possibility that SidC/SdcA may be activated by, or interact in complex with another bacterial or host cell protein during infection in order to directly catalyze cross-family small GTPase ubiquitination.

The connection between small GTPase ubiquitination and the activity of SidC/SdcA suggests that GTPase ubiquitination likely promotes *L.p.* pathogenesis and may play a role in SidC/SdcA-associated phenotypes such as ER-membrane recruitment and vacuolar expansion. Additionally, the cross-family ubiquitination of small GTPases with such disparate signaling roles within a common interface suggests that cross-family small GTPase ubiquitination targets an aspect of GTPase function common to all small GTPases, such as membrane association, nucleotide binding/hydrolysis, or structure of the Switch regions. Given these parameters, we propose two key models regarding the consequences of small GTPase ubiquitination during infection (**Fig 7C – Panel III**).

One potential model is that ubiquitination inhibits endogenous small GTPase signaling to prevent signaling detrimental to *L.p.* pathogenesis from occuring at the LCV membrane. We favor this model as *L.p.* is known to block LCV-membrane localized GTPases from their canonical binding partners using PTMs like AMPylation (Müller *et al*., 2010), phosphocholination (Mukherjee *et al*., 2011), or through the secretion of effectors like LidA, which can bind multiple Rabs at several orders of magnitude more strongly than mammalian Rab binding proteins (Schoebel *et al*., 2011). The use of ubiquitination as an additional mechanism to attenuate the binding of LCV-localized GTPases to their cognate signaling partners would be consistent with the significant degree of functional redundancy within the *L.p.* effector arsenal. Given the link between SidC/SdcA and membrane recruitment to the LCV (Luo and Isberg, 2004; Ragaz *et al*., 2008; Hsu *et al*., 2014), we also find this model to be enticing, as it suggests that *L.p.* may have evolved a strategy to couple the membrane recruitment necessary for LCV expansion to the inactivation of small GTPases on recruited membranes.

The second model, which is not mutually exclusive from the first, is that ubiquitination may prevent membrane dissociation of GTPases from the LCV. This model is supported by recent work in which the secreted *L.p.* deubiquitinase Lem27/LotC was shown to deubiquitinate Rab10 and decrease Rab10 association with the LCV during infection (Liu *et al*., 2020). Additionally, the presence of ubiquitination in the HVD, which contains CAAX motifs, lends support to the idea that ubiquitination directly affects membrane association. By this model, ubiquitin may directly interact with small GTPase lipid tails or the LCV membrane to promote membrane association, for example, or may indirectly promote membrane association by inhibiting GAP-mediated GTP hydrolysis.

Altogether, our work characterizes an unprecedented phenomenon of cross-family small GTPase ubiquitination that occurs during *L.p.* infection. We determine that most small GTPase ubiquitination falls within a distinct C-terminal interface, and that ubiquitination requires recruitment to the LCV membrane. This study places the secreted effectors SidC and SdcA, which have key roles in promoting timely LCV maturation and *L.p.* pathogenesis, as playing a central but indirect role in GTPase ubiquitination. Our work positions *L.p.* as a tool to understand a small GTPase regulation in both infected and uninfected contexts. Further examination of the host and bacterial proteins required for cross-family small GTPase ubiquitination is warranted, as the mechanistic details of this phenomenon will provide insight into both eukaryotic regulation of small GTPase activity and bacterial strategies of host cell manipulation.

## Materials and methods

### Cell lines

HEK293T cells (female), HEK293 cells (female) stably expressing the Fcγ receptor IIb (HEK293 FcγR cells), and HeLa FcγR cells were cultured in Dulbecco’s Modified Eagle’s Medium (DMEM, GIBCO) containing 10% fetal bovine serum (FBS, VWR) at 37°C and 5% CO_2_. FcγR expressing cell lines were gifts from the lab of Dr. Craig Roy at Yale University. U937 cells (a gift from Dr. Michael Bassik at Stanford University) were cultured in RPMI-1640 (Corning) supplemented with 10% heat-inactivated FBS (VWR). U937 were differentiated into macrophage-like cells in 20 ng/mL phorbol 12-myristate 13-acetate (PMA, Sigma) for 72 hours, then re-plated in media without PMA and allowed to rest for 48 hours before *L.p* infection.

### Bacterial strains and plasmids

Experiments were performed with *Legionella pneumophila* serogroup 1, strain Lp01 or Lp02. Avirulent T4SS-null strains were derived as previously described (Berger and Isberg, 1993; Berger *et al*., 1994). *L. pneumophila* strains were grown on Charcoal Yeast Extract (CYE) agar plates or AYE broth supplemented with (FeNO_3_ 0.135g/10mL) and cysteine (0.4g/10mL). Growth media for Lp02 thymidine auxotroph-derived strains was supplemented with 100 ug/mL thymidine. For strains carrying complementation plasmids, chloramphenicol (5 µg/mL) was supplemented for plasmid maintenance, and IPTG (1 mM) was added for 2 hours of induction prior to infection. The unmarked gene deletion Δ*sidC-sdcA* and Δ*drrA* strains were derived from the parental strain using allelic exchange as described previously (Berger *et al*., 1994). Rab5A, Rab5B, and Rab5C coding sequences were amplified from HeLa cDNA and cloned into a pcDNA3.1 mammalian expression vector containing the appropriate N-terminal tag (3XFlag or mCherry). Rab5A, Rab1A, and Rab10 CAAX deletion inserts were derived from appropriate full-length plasmid by PCR amplification of the desired region.

### Infection of cultured mammalian cells with *L.p*

Infections with *L.p.* were performed as previously described (Treacy-Abarca and Mukherjee, 2015). *L.p.* heavy patches grown for 48 h on CYE plates were either used directly for infection, or for overnight liquid cultures in AYE medium until reaching an OD600 of ∼3. *L.p.* from the overnight culture was enumerated and the appropriate amount was opsonized with *L.p*.-specific antibodies at a dilution of 1:2000 in cell growth medium for 20 min. HEK293 FcγR were grown on poly-lysine coated cell culture plates to a confluency of 80% and infected with the *L.p.* WT strain or the isogenic Δ*dotA* mutant strain at a multiplicity of infection (MOI) of 1-100 as indicated. The infection was synchronized by centrifugation of the plates at 1000xg for 5 min. To prevent internalization of any remaining extracellular bacteria at later timepoints, cells were washed three times with warm PBS after 1 h of infection and fresh growth medium was added. Cells were collected for down-stream processing at the indicated timepoints. Uninfected samples used as controls for infection experiments were mock-infected using media and opsonization antibody only.

### Sample preparation for proteomics analysis

HEK293 FcγR infected for 1 h or 8 h with the *L.p.* WT strain Lp01 or the isogenic *ΔdotA* mutant were infected at an MOI of 100. Uninfected HEK293 FcγR cells were included as a control. Cells were washed with ice-cold PBS, collected and the pellet was frozen at –80°C. Cell pellets were lysed by probe sonication in three pulses of 20% amplitude for 15 s in a lysis buffer consisting of: 8 M urea, 150 mM NaCl, 100 mM ammonium bicarbonate, pH 8; added per 10 ml of buffer: 1 tablet of Roche mini-complete protease inhibitor EDTA free and 1 tablet of Roche PhosSTOP. In order to remove insoluble precipitate, lysates were centrifuged at 16,100 g at 4°C for 30 min. A Bradford Assay (Thermo) was performed to measure protein concentration in cell lysate supernatants. 6 mg of each clarified lysate was reduced with 4 mM tris(2-carboxyethyl)phosphine for 30 min at room temperature and alkylated with 10 mM iodoacetamide for 30 min at room temperature in the dark. Remaining alkylated agent was quenched with 10 mM 1,4-dithiothreitol for 30 min at room temperature in the dark. The samples were diluted with three starting volumes of 100 mM ammonium bicarbonate, pH 8.0, to reduce the urea concentration to 2 M. Samples were incubated with 50 μg of sequencing grade modified trypsin (Promega) and incubated at room temperature with rotation for 18 hr. The sample pH was reduced to approximately 2.0 by the addition of 10% trifluoroacetic acid (TFA) to a final concentration of 0.3% trifluoroacetic acid. Insoluble material was removed by centrifugation at 16,000 g for 10 min. Peptides were desalted using SepPak C18 solid-phase extraction cartridges (Waters). The columns were activated with 1 ml of 80% acetonitrile (I), 0.1% TFA, and equilibrated 3 times with 1 ml of 0.1% TFA. Peptide samples were applied to the columns, and the columns were washed 3 times with 1 ml of 0.1% TFA. Peptides were eluted with 1.2 ml of 50% I, 0.25% formic acid. Peptides were divided for global protein analysis (10 μg) or diGly-enrichment (remaining sample), and lyophilized.

### diGlycine peptide enrichment by immunoprecipitation

Peptide samples were subjected to ubiquitin remnant immunoaffinity. 10 uL of PTMScan® Ubiquitin Remnant Motif (K-ε-GG) Antibody Bead Conjugate purification (Cell Signaling) slurry was used per 1 mg peptide sample. Ubiquitin remnant beads were washed twice with IAP buffer, then split into individual 1.7 mL low bind tubes (Eppendorf) for binding with peptides. Peptides were dried with a centrifugal evaporator for 12 hours to remove TFA in the elution. The lyophilized peptides were resuspended in 1 ml of IAP buffer (50 mM 4-morpholinepropnesulfonic acid, 10 mM disodium hydrogen phosphate, 50 mM sodium chloride, pH 7.5). Peptides were sonicated and centrifuged for 5 minutes at 16,100g. The soluble peptide supernatant was incubated with the beads at 4°C for 90 minutes with rotation. Unbound peptides were separated from the beads after centrifugation at 700g for 60 seconds. Beads containing peptides with di-glycine remnants were washed twice with 500 µL of IAP buffer, then washed twice with 500 µL of water, with a 700g 60s centrifugation to allow the collection of each wash step. Peptides were eluted twice with 60 µL of 0.15% TFA. Di-glycine remnant peptides were desalted with UltraMicroSpin C18 column (The Nest Group). Desalted peptides were dried with a centrifugal adaptor and stored at –20°C until analysis by liquid chromatograph and mass spectrometry.

### Mass spectrometry data acquisition and processing

Samples were resuspended in 4% formic acid, 4% acetonitrile solution, separated by a reversed-phase gradient over a nanoflow column (360 µm O.D. x 75 µm I.D.) packed with 25 cm of 1.8 µm Reprosil C18 particles with (Dr. Maisch), and directly injected into an Orbitrap Fusion Lumos Tribrid Mass Spectrometer (Thermo). Total acquisition times were 120 min for protein abundance, 100 min for phosphorylation, and 70 min for ubiquitylation analyses. Specific data acquisition settings are detailed in **Supplemental Table 1**. Raw MS data were searched with MaxQuant against both the human proteome (UniProt canonical protein sequences downloaded January 11, 2016) and the *Legionella Pneumophila Philadelphia* proteome (downloaded July 17, 2017). Peptides, proteins, and PTMs were filtered to 1% false discovery rate in MaxQuant (Cox *et al*., 2014). Principal Component analysis of normalized MS Intensities of experimental conditions (control, Δ*dotA*-1h, Δ*dotA*-8h, WT-1h, WT-8h) was performed using the factoextra R package as implemented by the artMS bioconductor package. The plot illustrates the relationship between the variables (conditions) and the principal components, where each variable is represented as a vector, and the direction and length of the vectors indicate how each variable contributes to the two principal components. If two vectors are close together indicates a strong positive correlation between those two variables, i.e. they contribute to the principal components in a similar way. Statistical analysis of quantifications obtained from MaxQuant was performed with the artMS Bioconductor package (version 0.9) (Jimenez-Morales *et al*., 2019). Each dataset (proteome and ubiquitinome) was analyzed independently. Quality control plots were generated using the artMS quality control functions. The site-specific relative quantification of posttranslational modifications required a preliminary step consisting of providing the ptm-site/peptide-specific annotation (“artmsProtein2SiteConversion()” function). artMS performs the relative quantification using the MSstats Bioconductor package (version 3.14.1) (Choi *et al*., 2014). Contaminants and decoy hits were removed. Samples were normalized across fractions by median-centering the Log_2_-transformed MS1 intensity distributions (**Fig 1-S1B, Fig 1-S3B**). **Imputation strategy:** Log2FC for protein/sites with missing values in one condition but found in ≥2 biological replicates of the other condition of any given comparison were estimated by imputing intensity values from the lowest observed MS1-intensity across sample peptides (Webb-Robertson *et al*., 2015); p-values were randomly assigned between 0.05 and 0.01 for illustration purposes.

### Subcellular compartment analysis, functional enrichment analysis, and small GTPase sequence alignment

Statistically significant changes were selected by applying the joint thresholds of |Log2FC| ≥ 1, adj.-p-value < 0.05. Imputed values were also considered significant and are indicated in figures separately from non-imputed values. WT1hr-Control, WT8hr-Control, Δ*dotA*1hr-Control, and Δ*dotA*8hr-Control comparisons were filtered using these significance criteria for subsequent analyses. Subcellular compartment analysis was performed by tabulating the number of significantly regulated proteins per compartment based on subcellular localization identifiers from UniProt. Biological pathway and protein complex enrichment was performed using Metascape (Zhou *et al*., 2019) (https://metascape.org). The following ontology sources were used for analysis: GO Biological Processes, KEGG Pathway, GO Molecular Functions, GO Cellular Components, Reactome Gene Sets, Hallmark Gene Sets, Canonical Pathways, BioCarta Gene Sets, CORUM, WikiPathways and PANTHER Pathway. Significant enrichment terms were selected using the combined thresholds of p-value < 0.01, a minimum count of 3 proteins, and an enrichment factor > 1.5. Proportional Venn diagrams were created using DeepVenn (Hulsen, 2022) and recolored in Adobe Illustrator. Proteins within the Ras superfamily were defined based on the “Ras small GTPase superfamily” definition in the HUGO Gene Nomenclature Committee database (https://www.genenames.org/, HGNC group ID = 358). Sequence alignment was performed using Jalview (Waterhouse *et al*., 2009).

### Cell lysis, immunoprecipitation, and immunoblot analysis

HEK293 FcγR cells grown on poly-lysine coated plates were treated as indicated, washed three times with ice-cold PBS and harvested with a cell scraper. Cells were pelleted at 3000xg for 10 minutes at 4C. For ubiquitin pulldown assays using the SignalSeeker kit (Cytoskeleton Inc), cells were lysed in provided BlastR buffer with protease inhibitor and NEM, and total protein concentration measured using Precision Red Advanced protein assay. Lysates were diluted to 1 mg/mL, and 1 mL of diluted lysate was incubated with either unconjugated (control) or ubiquitin binding domain conjugated beads for 2 hours at 4C on a rotating platform. Beads were washed three times in wash buffer, and bound proteins were eluted using kit spin columns. For all other immunoblots, cell pellets were resuspended in RIPA buffer supplemented with cOmplete Protease Inhibitor Cocktail (Roche), phenymethylsulphonyl fluoride (PMSF, 1 mM), and 10 mM NEM and lysed under constant agitation for 20 min at 4°C. Cell debris was removed by centrifugation at 16,000xg for 20 min at 4°C. Protein concentration was measured using the Pierce 660nm Protein Assay Kit or the Pierce™ BCA Protein Assay Kit (Thermo Fisher Scientific). For each sample, 20-30 μg of proteins were denatured in SDS sample buffer/5% β-mercaptoethanol at 95°C for 5 min, loaded on 8-12% SDS-polyacrylamide gels and separated by SDS-PAGE. Proteins were transferred to PVDF membranes (0.45 μm, Millipore) at 30 V, 4°C for 16 h. For total ubiquitin blots (**Fig 1D**), total protein was quantified before blocking using Invitrogen No-Stain Protein Labeling Reagent. Membranes were washed with PBS-T (PBS/ 0.1% Tween-20 (Thermo Fisher Scientific)), blocked with 5% Blotting Grade Blocker Non Fat Dry Milk (Bio-Rad) for 1 h at room temperature and incubated with the primary antibodies diluted in blocking buffer/0.02% (w/v) sodium azide overnight at 4°C. Membranes were washed three times with PBS-T and incubated with Goat Anti-Mouse IgG (H+L) HRP Conjugate (Thermo Fisher Scientific), Goat Anti-Rabbit IgG (H+L) (Thermo Fisher Scientific), HRP Conjugate, diluted at 1:5000 in blocking buffer for 60 min at room temperature. After three washes with PBS-T, membranes were incubated with Amersham ECL Western Blotting Detection Reagent (Global Life Science Solutions) for 1 min and imaged on a ChemiDoc Imaging System (BioRad).

### Immunoblot quantification

Images were exported from ImageLab (BioRad) as 16-bit tiff and analyzed in ImageJ. Plot profiles were generated for each lane and the integrated density was calculated using the ImageJ built in gel analyzer tools. Total ubiquitin signal was normalized to total protein, and the fold change was calculated compared to the appropriate uninfected control. To calculate normalized Rab monoubiquitination intensity, integrated density was measured for the unmodified band at sub-saturated exposure. Integrated density was measured for the higher molecular weight monoubiquitination band at the lowest exposure in which this band was visible. Normalized Rab monoubiquitination was calculated as follows: IntDen monoUb/(IntDen monoUb + InDen unmod Ub). To standardize these values across biological replicates, values are represented as a percentage of the WT infection condition for each replicate.

### Immunofluorescence and image analysis

HeLa FcγR cells were grown on poly-lysine coated coverslips in 24well cell culture plates. Cells were treated as indicated, washed three times with PBS and fixed in 4% paraformaldehyde/PBS for 15 min at room temperature. Cells were then treated with 2% BSA, 0.5% saponin in PBS (blocking/permeabilization buffer) for 1h at RT. Cells were stained with primary antibodies diluted in blocking/permeabilization buffer overnight at 4C, washed three times with PBS and stained with secondary antibodies diluted in blocking/permeabilization buffer for 1h at RT. Cells were then stained with Hoechst33342 at 1:2000 in PBS for 10 min and washed three times with PBS. Coverslips were dipped three times into purified ddH_2_O to remove salts, dried and mounted on microscopy glass slides with Prolong Diamond antifade 5 (Thermo Fisher Scientific). Slides were cured overnight at room temperature and imaged on a spinning disk Eclipse Ti2-E inverted microscope (Nikon). Images were analyzed in Fiji. Experimental conditions were blinded either before image acquisition, or before image analysis using the Fiji Blind Analysis Tools plugin filename encrypter. For LCV scoring, max intensity Z projections were generated. LCVs were scored positive if the LCV region was visible in the protein marker of interest channel only (i.e. without the *L.p.* marker). All LCV area measurements were carried out in Fiji using the freehand selection tool.

### Cell transfections

All transfections were performed with jetPRIME (Polyplus). HEK293 FcγR or HeLa FcγR cells were grown to 60% confluency and transfected according to the manufacturer’s recommendations. For transfection of plasmid DNA, 0.25 μg DNA was used for 24well plates, 1-2 μg DNA for 6 well plates, and 2-3 ug for 60 mm plates. 24h after transfection, cells were treated as indicated and analyzed or harvested.

## Data availability

The mass spectrometry data files have been deposited to the ProteomeXchange Consortium (http://proteomecentral.proteomexchange.org) via the PRIDE partner repository with the dataset identifier PXD019217 (Vizcaíno *et al*., 2016).

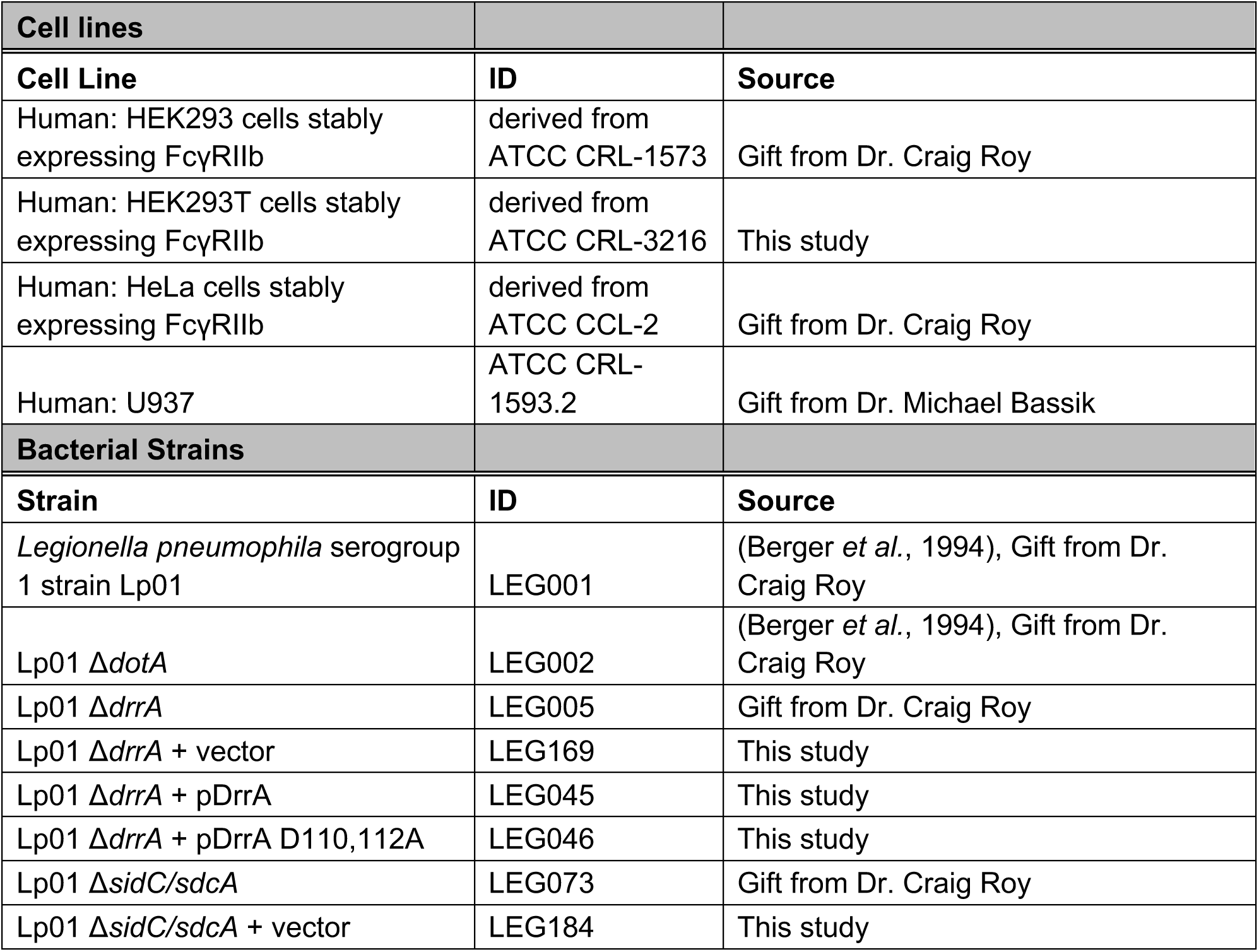
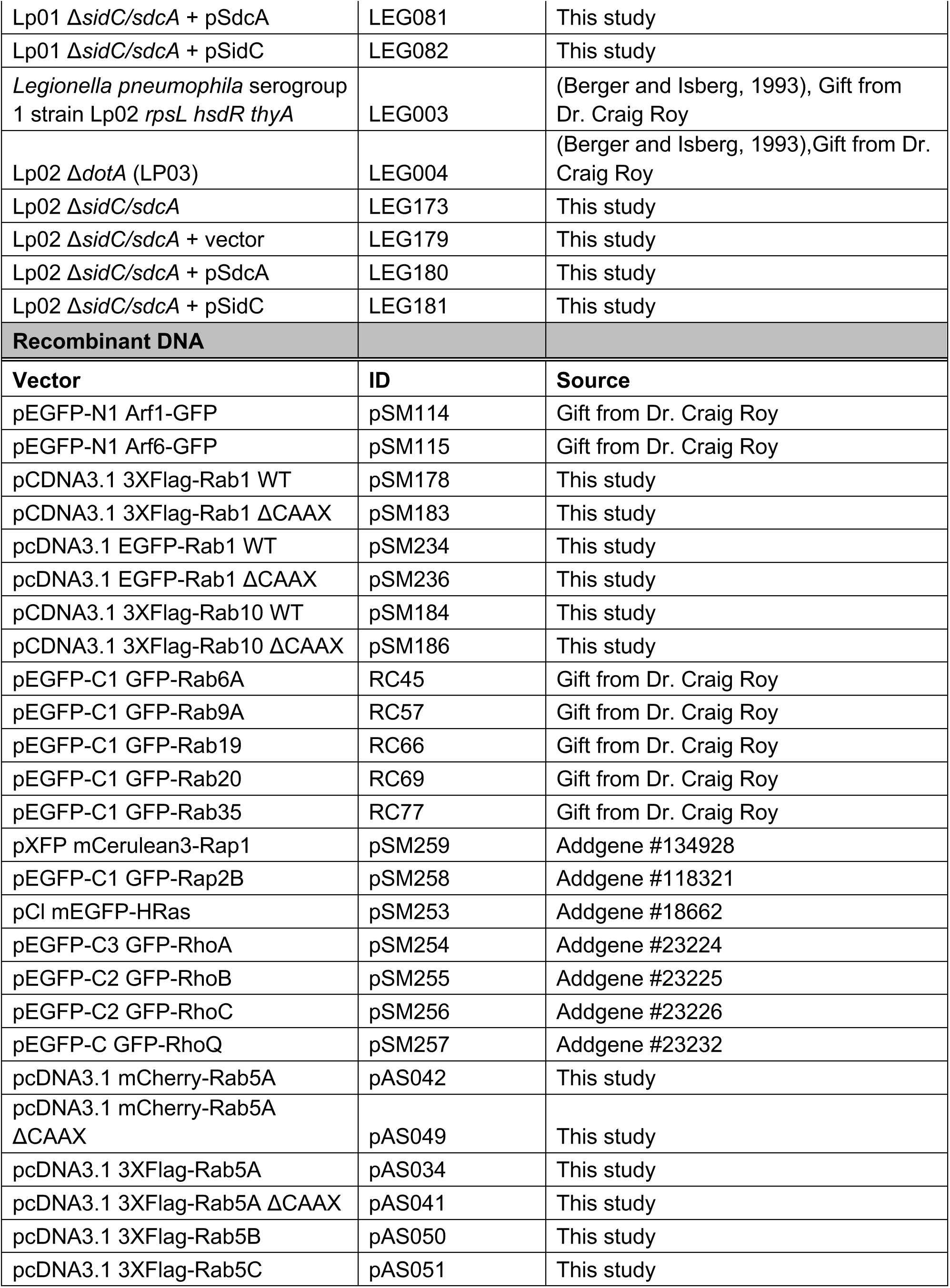
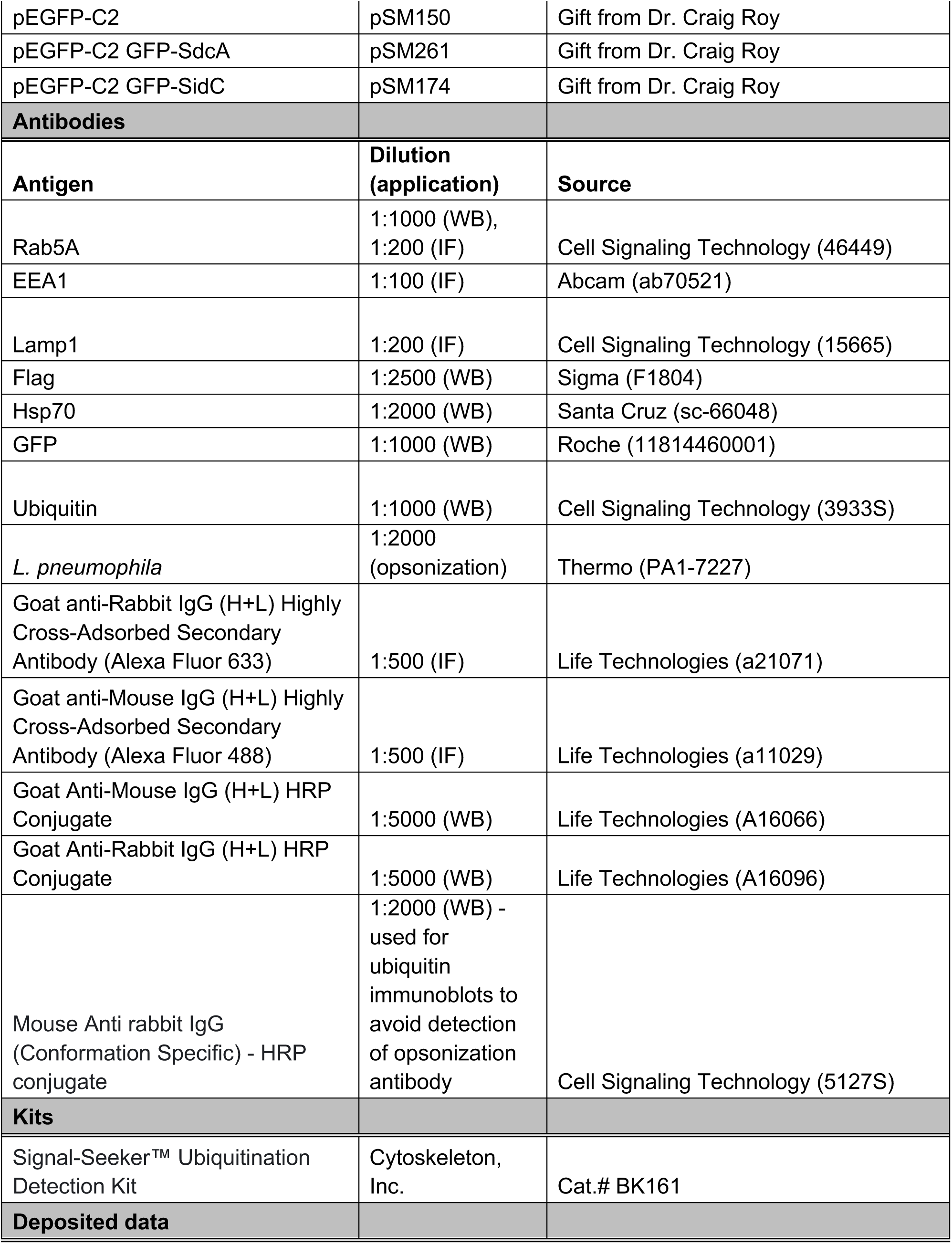
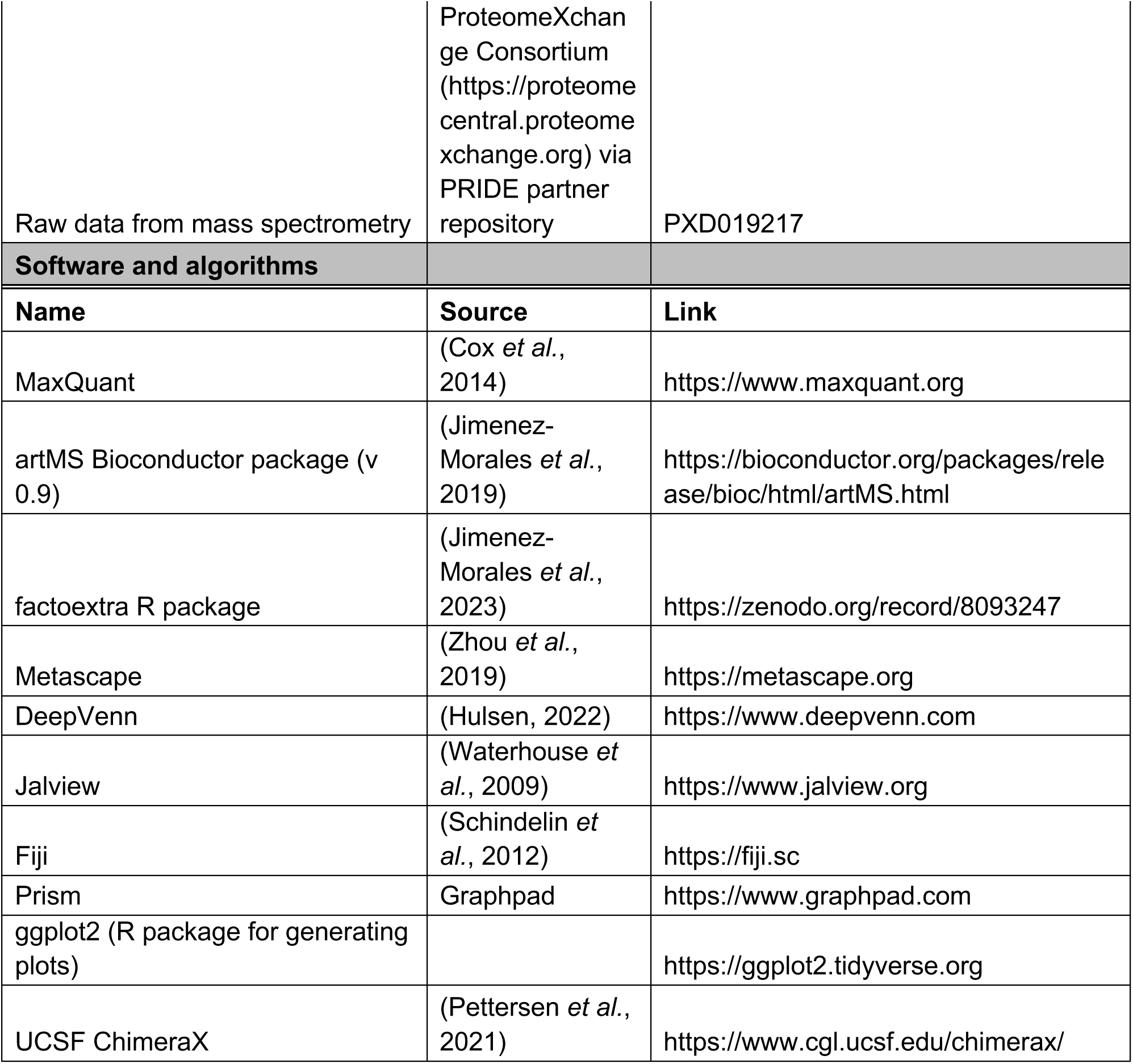
Key resources table.

## Supplemental Information

**Figure 1-S1:**
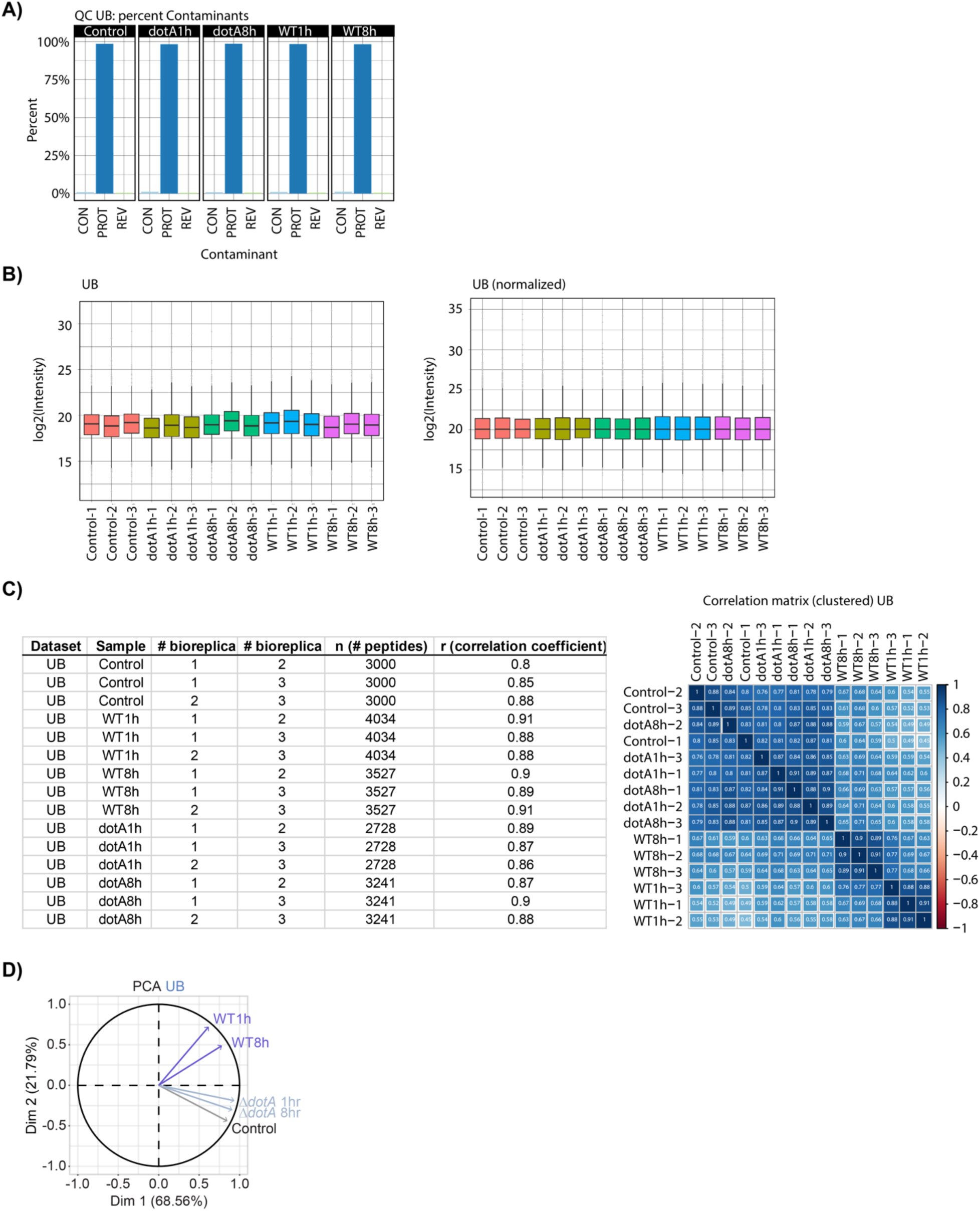
Quantification and quality control plots of ubiquitinomics data. Related to Figure 1. Quality control plots for diGly enriched dataset were generated using the artMS Bioconductor package (version 0.9) (Jimenez-Morales *et al*., 2019). **(A**) Percent of contaminants (CON), proteins (PROT) and reversed sequences (REV) in each experimental condition (control, dotA-1h, dotA-8h, WT-1h, WT-8h) were quantified to adjust the false-discovery-rate (FDR). **(B**) Samples were normalized across fractions by median-centering the Log2-transformed MS1 intensity distributions. **(C**) Correlation table and matrix showing the clustering of the different experimental conditions. **(D**) Principal Component analysis of normalized MS Intensities of experimental conditions (control, ΔdotA-1h, ΔdotA-8h, WT-1h, WT-8h). PC1 and PC2 captured most of the variability. Loading variables are represented as vectors. The smaller angle between control and the mutant time points (Δ*dotA*-1h, Δ*dotA*-8h) implies a larger positive correlation between them, as opposed to a lower correlation (larger angle) between the Control and the WT strain.

**Figure 1-S2:**
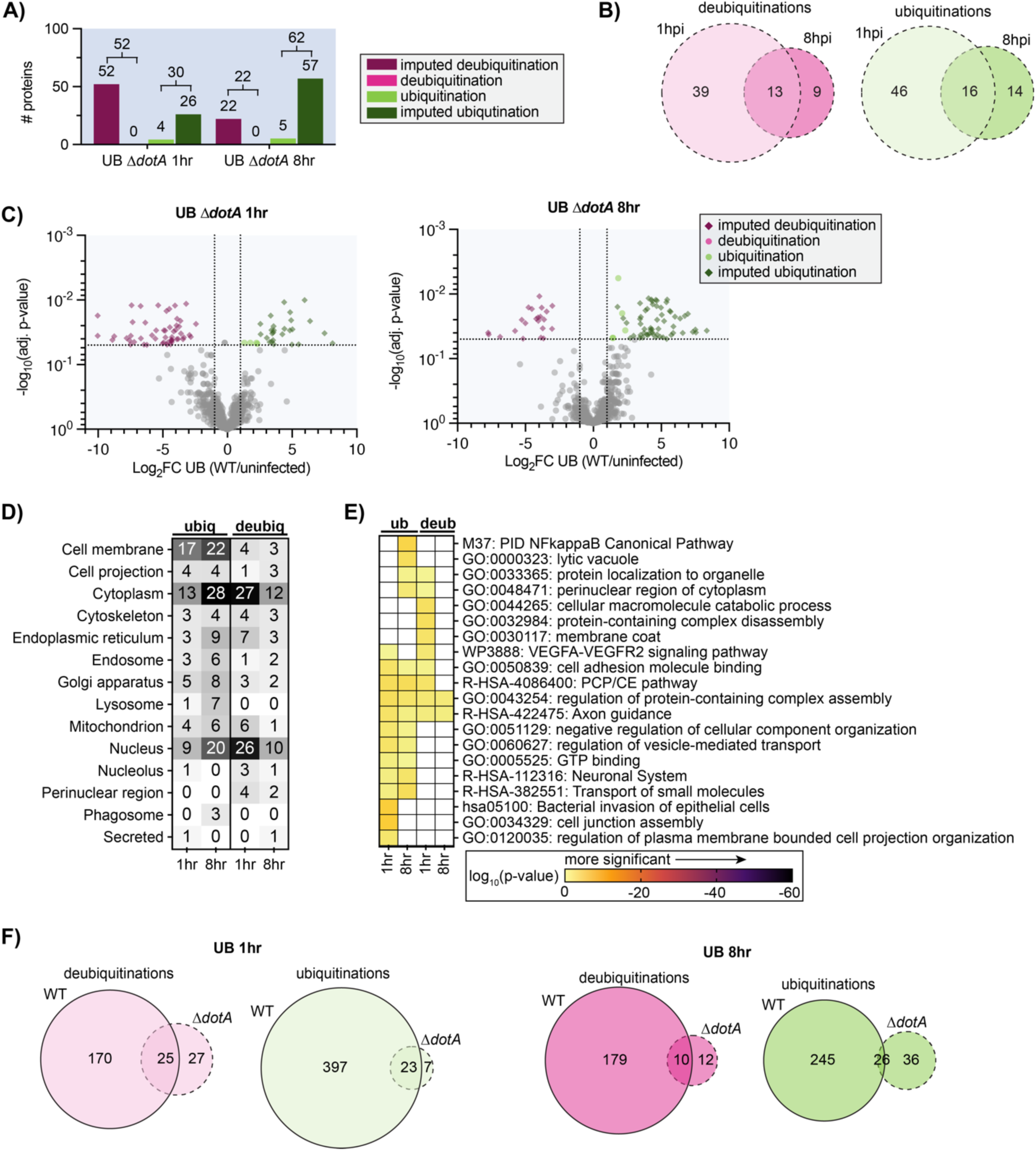
Analysis of T4SS-independent (*L.p.* Δ*dotA*) changes in the host cell ubiquitinome. Related to Figure 1. (**A**) Counts of proteins with a significant increase (green) or decrease (magenta) in ubiquitination compared to uninfected control for Δ*dotA* 1hr and Δ*dotA* 8hr (same data shown in Figure 1B, repeated here for clarity). **(B**) Overlap of proteins with a significant increase (green) or decrease (magenta) in ubiquitination compared to uninfected control in the 1-hour vs 8-hour *L.p*. Δ*dotA* infected conditions. **(C**) Volcano plot representation of all ubiquitinome data in Δ*dotA* vs uninfected comparison at 1-and 8-hours post-infection. Imputed values are shown as diamonds. Significance threshold is indicated by the dotted line. **(D**) Subcellular localization analysis of proteins with a significant increase or decrease in ubiquitination compared to uninfected control during *L.p*. Δ*dotA* infection for 1 or 8 hours. **(E**) Metascape pathway and protein complex analysis of proteins with a significant increase or decrease in ubiquitination compared to uninfected control during *L.p*. Δ*dotA* infection for 1 or 8 hours. Terms not significantly enriched for a given experimental condition are represented by white boxes. **(F**) Overlap of proteins with a significant increase (green) or decrease (magenta) in ubiquitination compared to uninfected control in the WT *L.p*. vs. Δ*dotA* infected conditions, at both 1– and 8-hours post-infection.

**Figure 1-S3:**
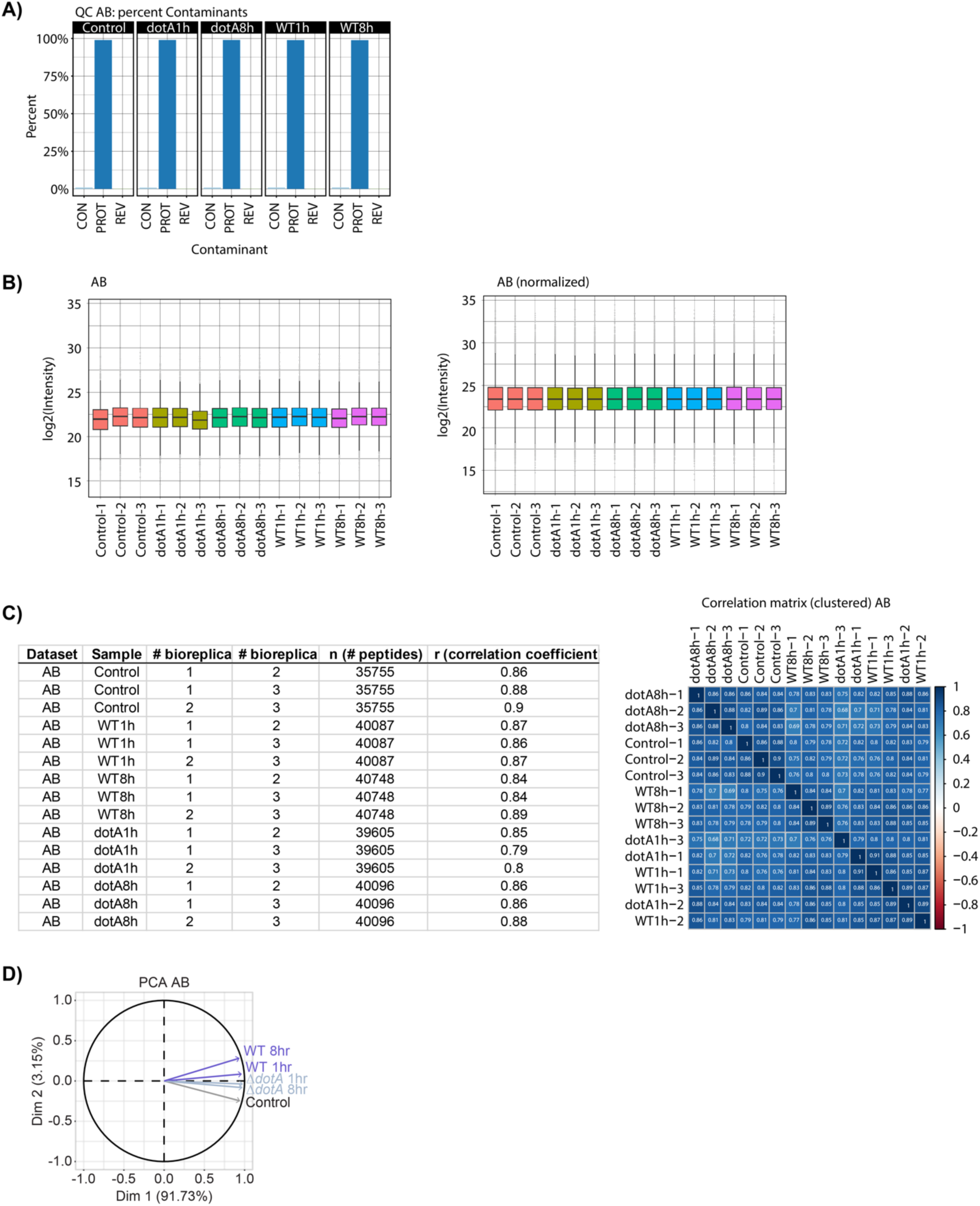
Quantification and quality control plots of abundance data. Related to Figure 1. Identical analysis as in Figure 1S1 using abundance data.

**Figure 1-S4:**
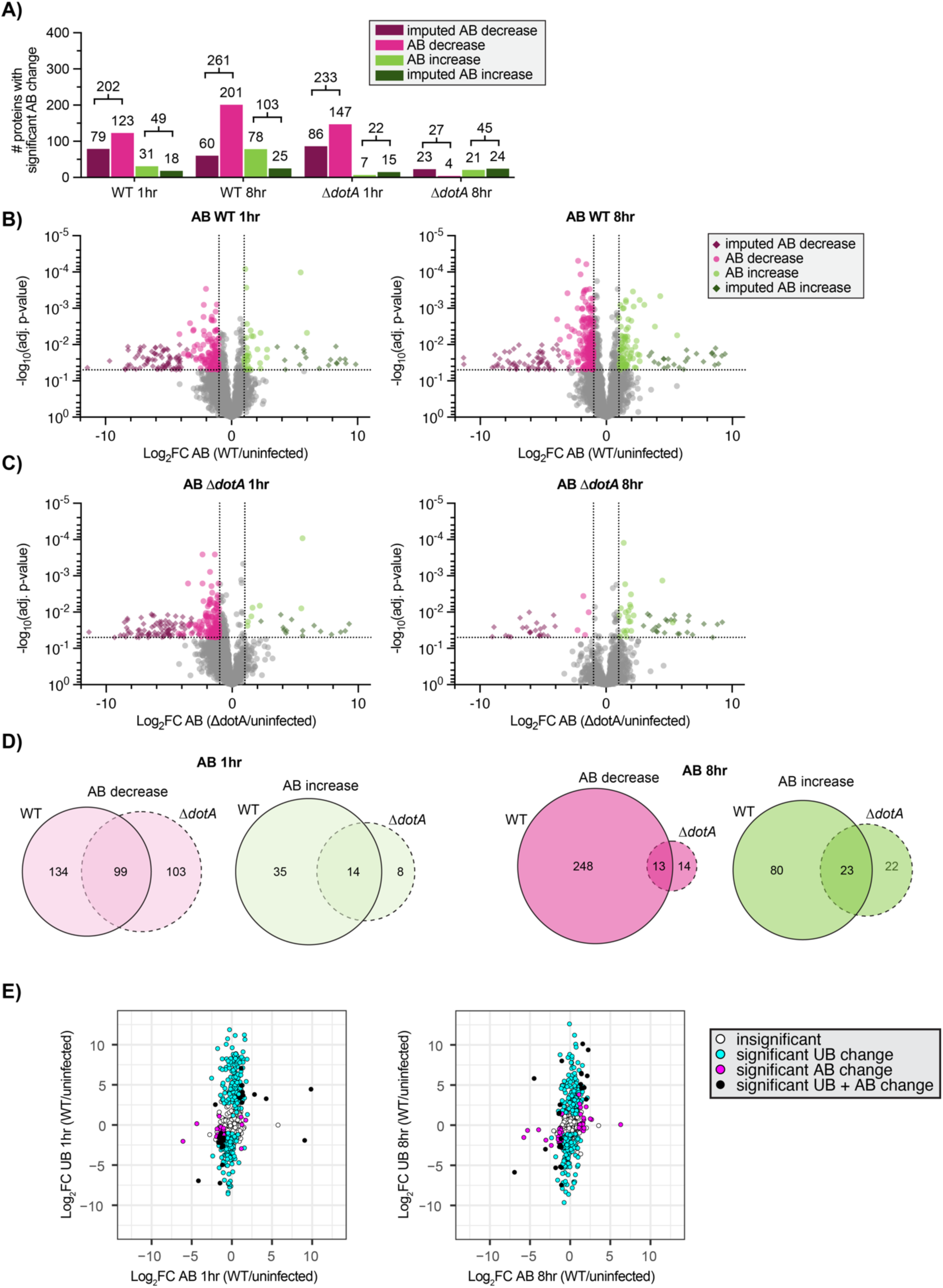
Analysis of *L.p.*-induced changes in host cell abundance. Related to Figure 1. Analysis of proteins with significant increases (green) or decreases (magenta) in abundance compared to uninfected control for WT and Δ*dotA* infected conditions, at both 1 and 8 hours post infection. **(A)-(D**) Comparable to analyses in Fig 1 and Fig 1S2. **(E**) Log2FC comparison of proteins with significant abundance change (grey points), significant ubiquitination change (light blue points), or both significant abundance and ubiquitination change (dark blue points) during infection with WT *L.p.* relative to uninfected control. Proteins with insignificant abundance and ubiquitination changes are shown in white. Data shown are at 1-hour post-infection (left) and 8-hours post-infection (right).

**Figure 2-S1:**
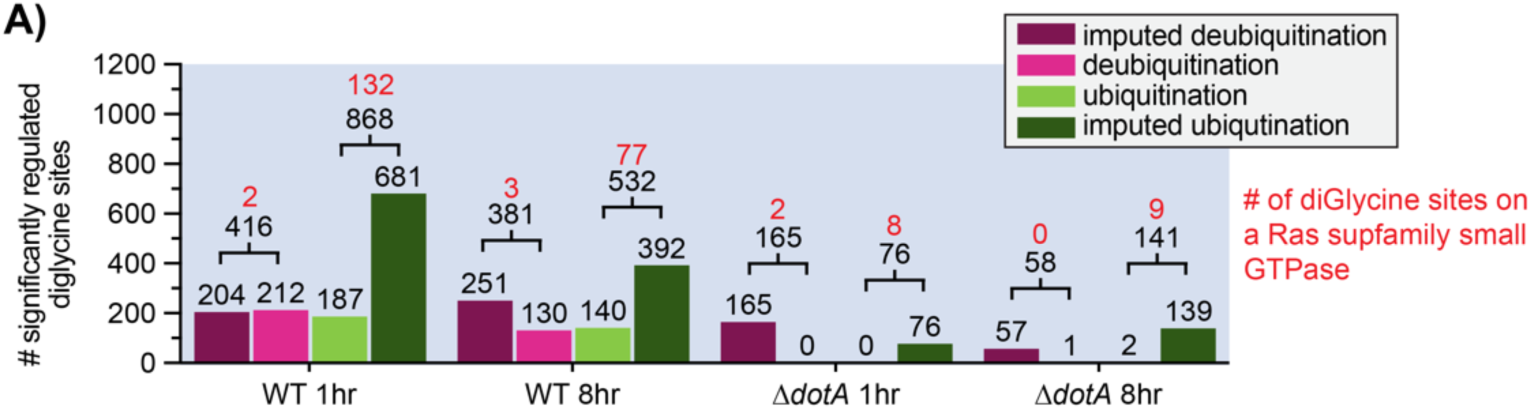
Quantification of diGlycine site data. Related to Figure 2. Counts of diGly sites with a significant increase (green) or decrease (magenta) compared to uninfected control for the indicated infection conditions. Significance threshold: |log2(FC)|>1, p<0.05. DiGlycine sites falling on small GTPases indicated in red.

**Figure 2-S2:**
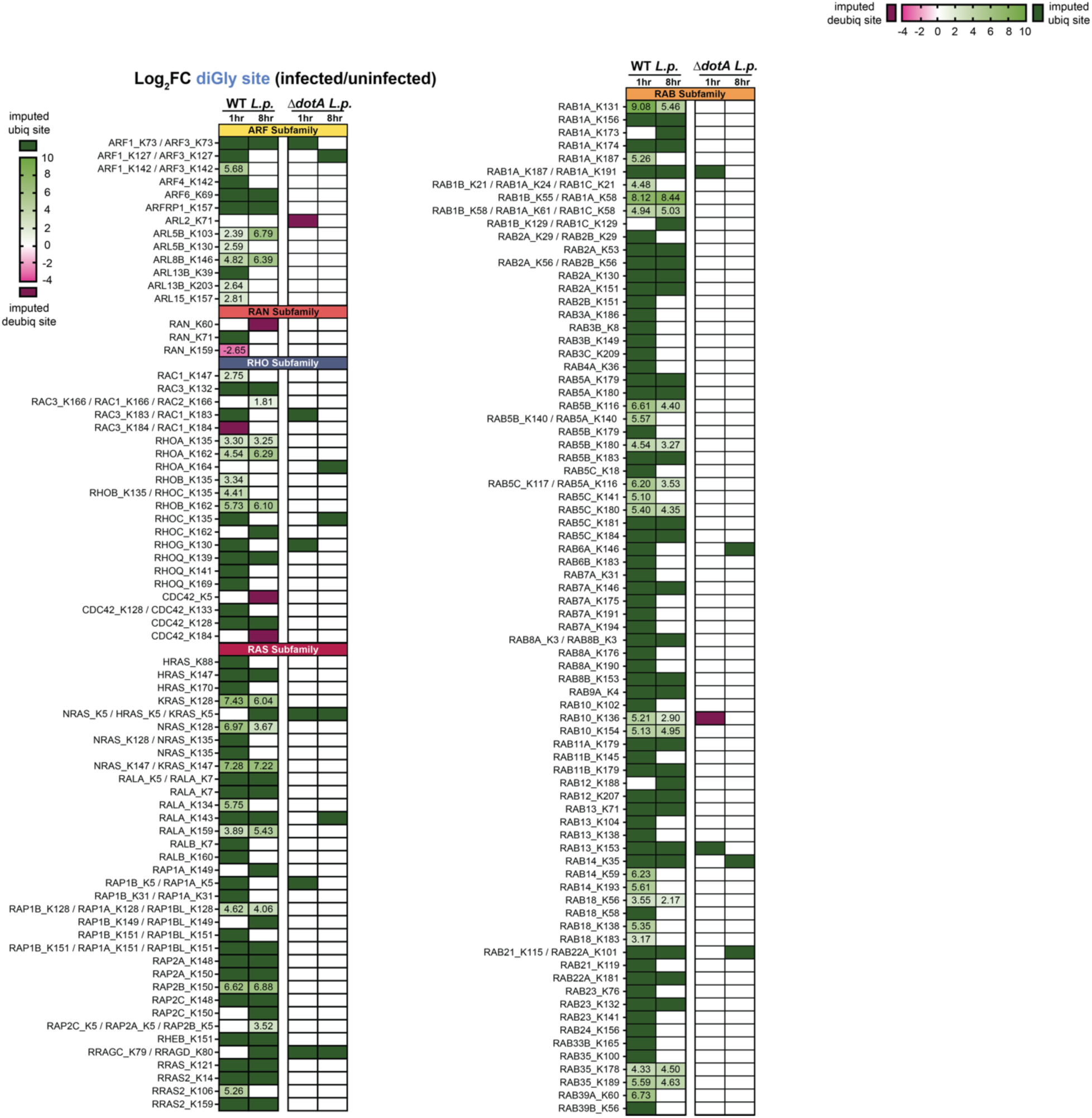
List of significantly regulated diGly sites falling on small GTPases. Related to Figure 2. All significantly changing diGlycine sites falling on small GTPases during WT and Δ*dotA L.p.* infection, organized by subfamily and infection timepoint. Cell color and number indicate Log2FC. White cells indicate no significant change. Due to the significant homology between various small GTPases, some trypsinized peptides could not be distinguished between multiple proteins. DiGlycine sites that could be assigned to multiple proteins are indicated with a “/”. In our downstream analyses, we use a conservative approach and only consider the first small GTPase listed (e.g. NRAS_K147 / KRAS_K147, only NRAS_K147 is counted as a small GTPase ubiquitination site).

**Figure 2-S3:**
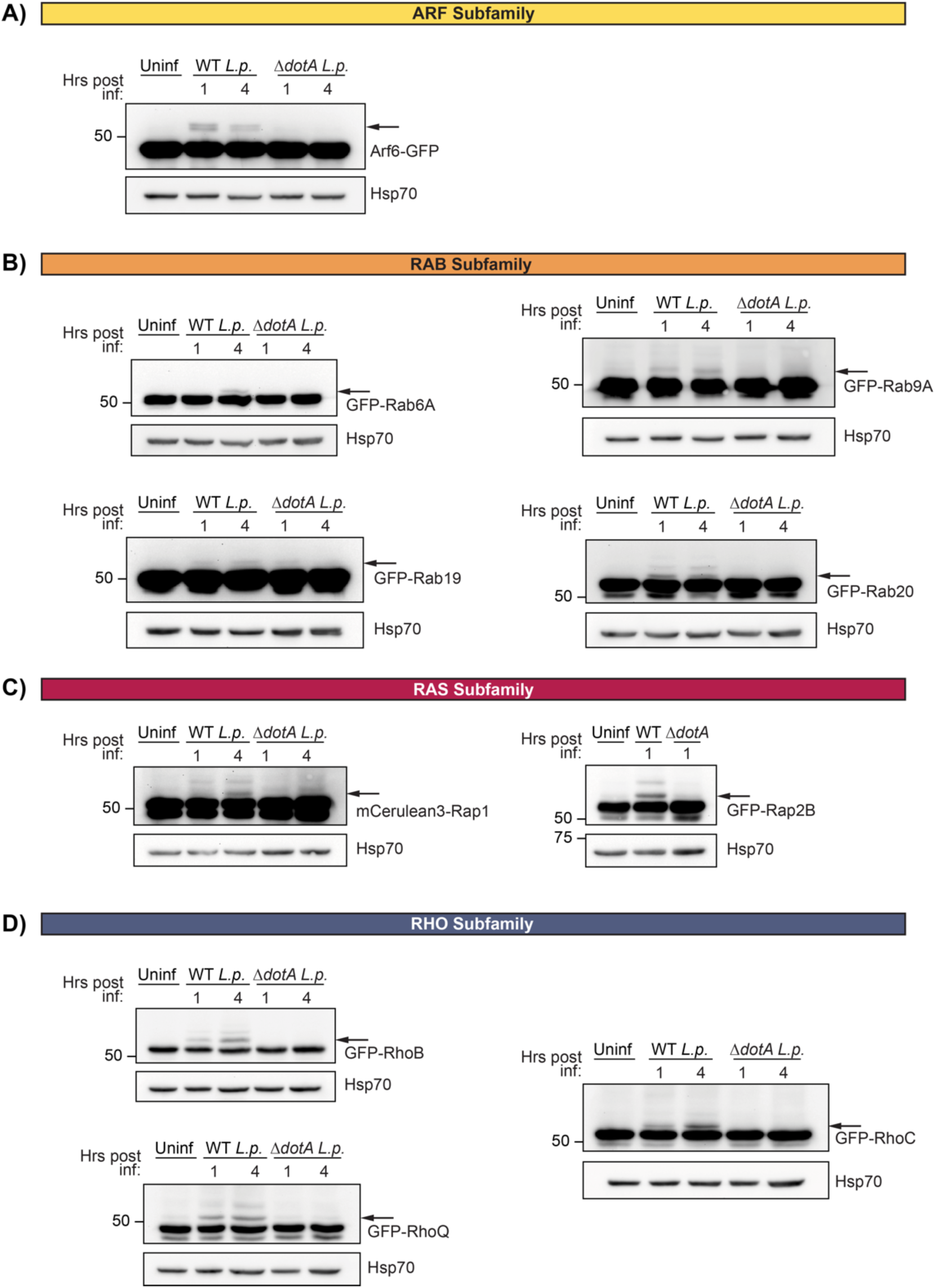
Extended immunoblot analysis of small GTPase ubiquitination confirmations. Related to Figure 2. Extension of Figure 2B-E. Immunoblot analysis of lysates prepared from HEK293T FcγR cells transiently transfected with the indicated GFP-tagged small GTPase, then infected with either WT or Δ*dotA L.p.* for 1 to 4 hours, or left uninfected. Blots were probed with anti-GFP and anti-Hsp70 antibodies. Monoubiquitinated GTPase indicated with an arrow.

**Figure 3-S1:**
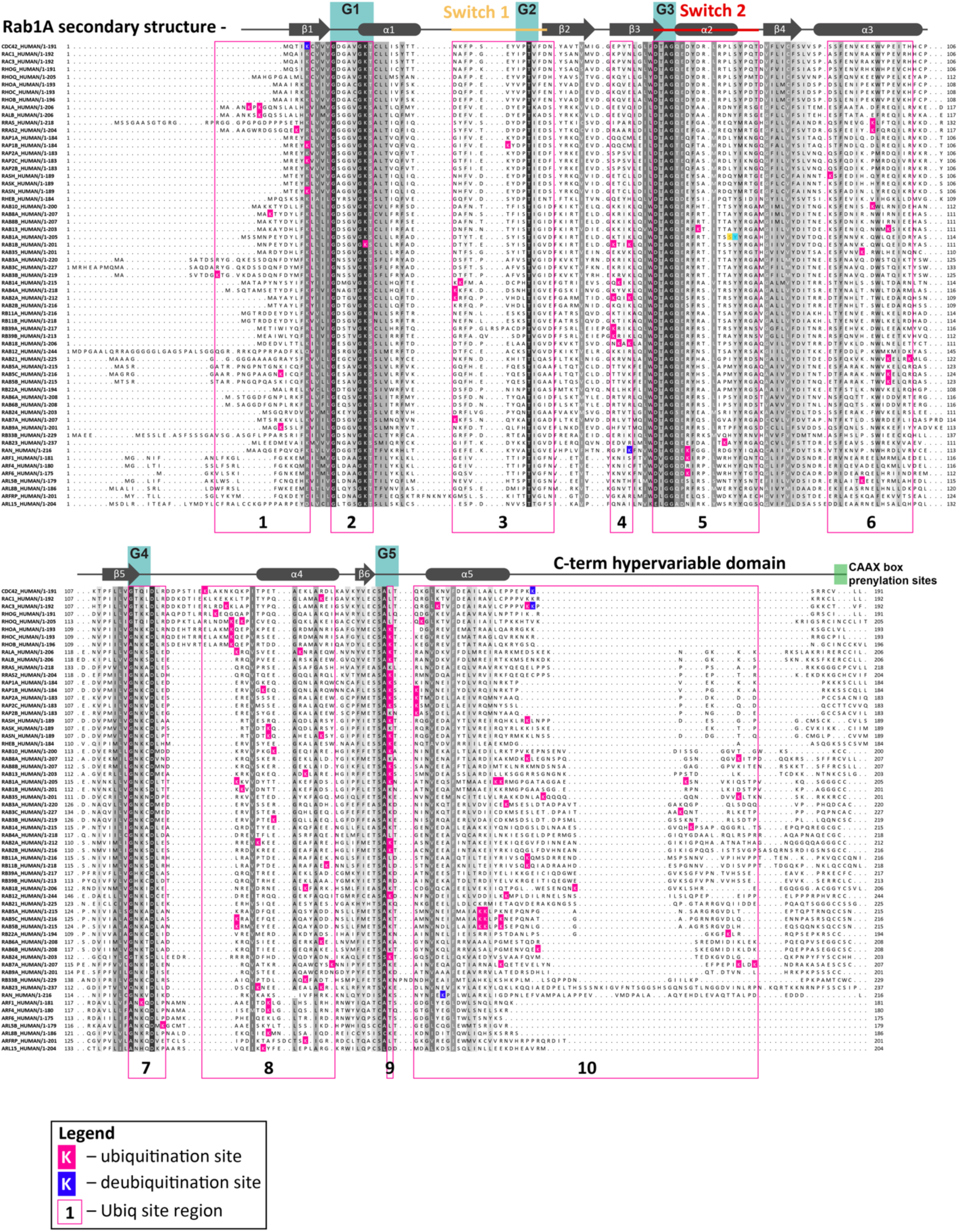
Sequence alignment of ubiquitinated small GTPases with UB sites indicated. Related to Figure 3. Sequence alignment of small GTPases ubiquitinated during infection. 61 of 63 ubiquitinated small GTPases represented, with RRAGC and ARL13B omitted for visual clarity due to their significant length. Sequence colored by conservation within the small GTPase superfamily, from white (non-conserved) to black (extremely highly conserved residue). Ubiquitinated residues at either 1– or 8-hours post infection are colored in bright pink and outlined in black, and deubiquitinated residues are colored in dark blue. Structural and functional regions indicated above, using Rab1A as an example. Regions frequently ubiquitinated across detected small GTPases are underlined in pink and numbered 1-10. ARL13B_K39 and RRAGC_K79 fall in UB region #2. ARL13B_K203 falls in UB region #10.

**Figure 4-S1:**
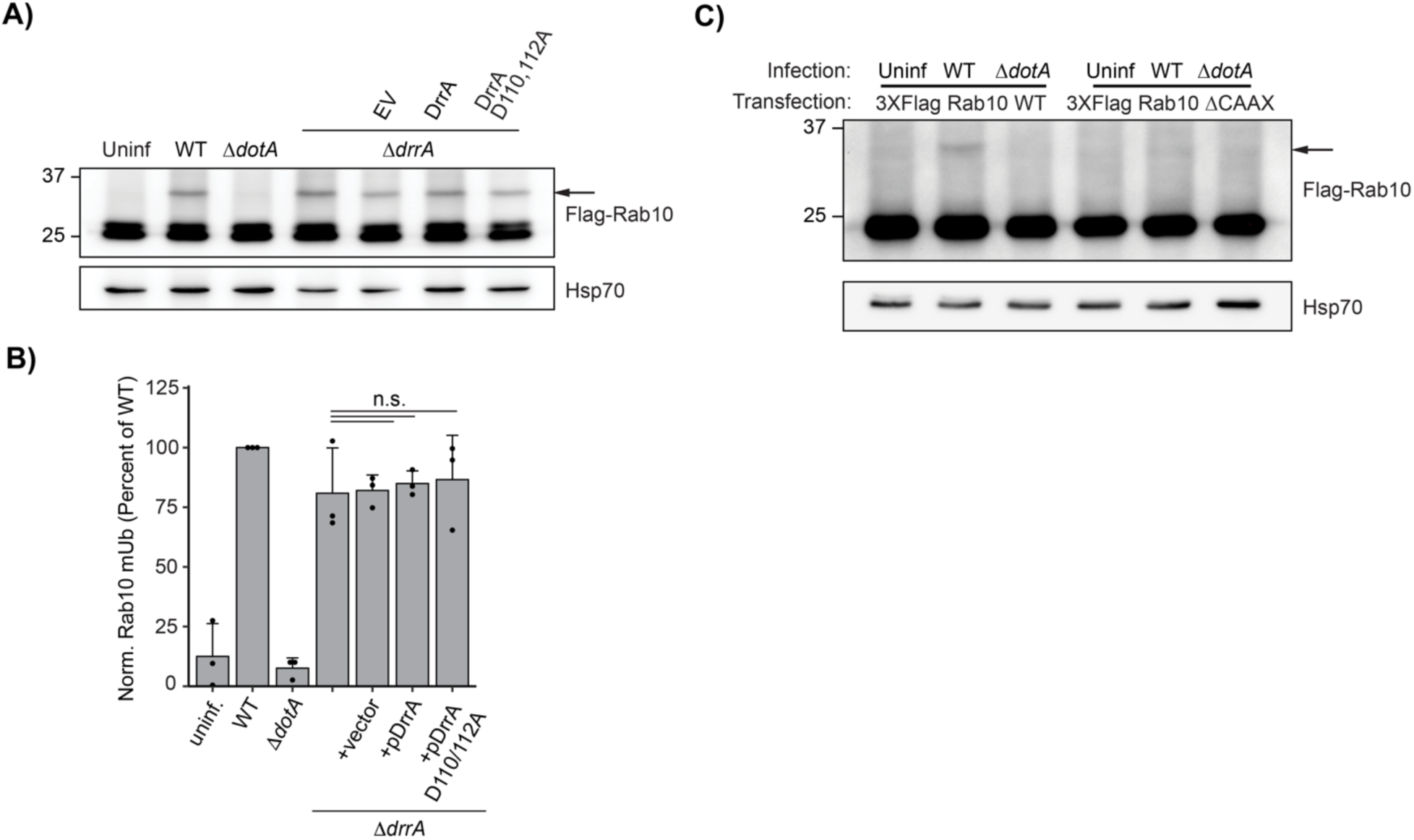
Rab10 ubiquitination is not controlled by DrrA and requires membrane association. (**A**) Immunoblot analysis of lysates prepared from HEK293T FcγR transfected with 3XFlag Rab10 and infected with a Δ*drrA L.p*. strain panel (WT, Δ*dotA*, Δ*drrA*, and Δ*drrA* complemented with empty vector or plasmid encoded DrrA WT or D110, 112A) for 1 hour (MOI=50). Monoubiquitinated Rab10 indicated with an arrow. **(B**) Quantification of biological replicates (N=3) of experiment shown in (A). Normalized Rab10 monoubiquitination intensity was calculated as a percentage of WT *L.p.* infection levels (see Methods). **(C**) Immunoblot analysis of monoubiquitination of Rab10 WT vs ΔCAAX during *L.p.* infection. HEK293T FcγR cells were transfected with either 3X Flag Rab10 WT or ΔCAAX, then infected with WT or Δ*dotA L.p.* for 1 hour (MOI=50) or left uninfected. Lysates were probed with anti-Flag antibody. Monoubiquitinated Rab10 indicated with an arrow.

**Figure 5-S1:**
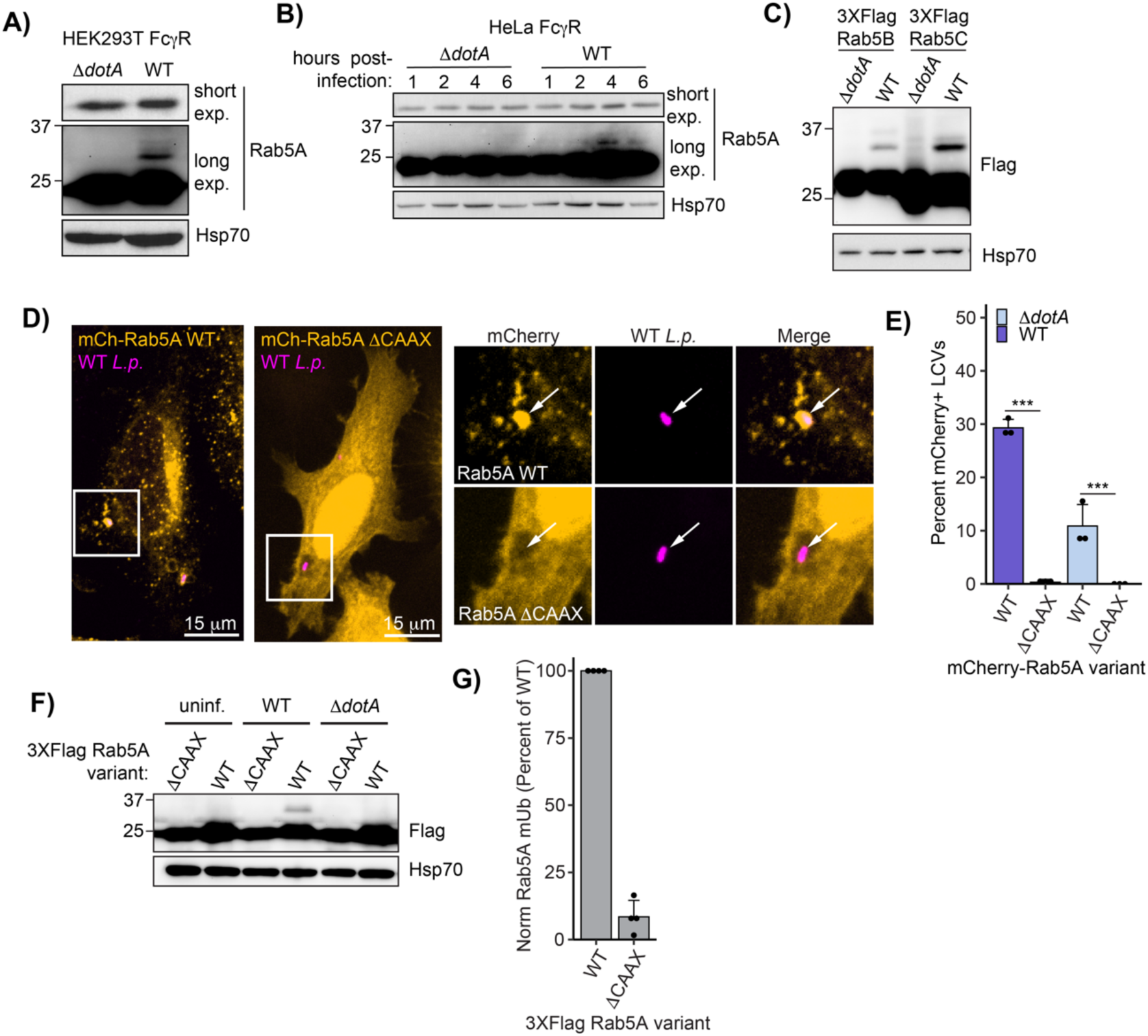
Rab5 ubiquitination reproduces across cell lines and isoforms and is dependent upon membrane localization. Related to Figure 5. (**A**) Immunoblot analysis of lysates prepared from HEK293T FcγR transfected with 3XFlag Rab5A and infected with *L.p.* WT or Δ*dotA*. **(B**) Immunoblot analysis of lysates prepared from HeLa FcγR transfected with 3XFlag Rab5A and infected with *L.p.* WT or Δ*dotA*. **(A**) Immunoblot analysis of lysates prepared from HEK293T FcγR transfected with 3XFlag Rab5B or 3XFlag Rab5C and infected with *L.p.* WT or Δ*dotA.* D) and **(E**) Immunofluorescence analysis of mCherry Rab5A WT or ΔCAAX LCV recruitment. HeLa FcγR cells were transfected with indicated construct, then infected for 1 hour with either WT or Δ*dotA L.p.*, fixed, and stained with anti-*Legionella* antibody. **(D**) Representative images, and **(E**) quantification of mCherry positive LCVs as in (D). **(F**) Immunoblot analysis of ubiquitination of Rab5A WT vs ΔCAAX during *L.p.* infection. HEK293T FcγR cells were transfected with either 3X Flag Rab5A WT or ΔCAAX, then infected with WT or Δ*dotA L.p.* for 4 hours or left uninfected. Lysates were probed with anti-Flag antibody. **(G**) Quantification of normalized Rab5A ΔCAAX mUb intensity as a percent of Rab5A WT mUb during WT *L.p.* infection (see Methods). Biological replicates (N=4) were carried out as in (G). *For all bar graphs:* bars represent mean value, error bars represent standard deviation. Individual points are values from each biological replicate. *Statistical analysis of LCV scoring quantification:* G test of independence was performed on pooled counts (positive vs. negative) from all biological replicates. Upon verifying significance (p<0.05), pairwise comparisons between strains were evaluated by post-hoc G-test using the Bonferroni-adjusted p-value as a significance threshold (p = 0.008). * = p<0.008, ** = p<0.0008, *** = p<0.00008, n.s. = p>0.008.

**Figure 7-S1:**
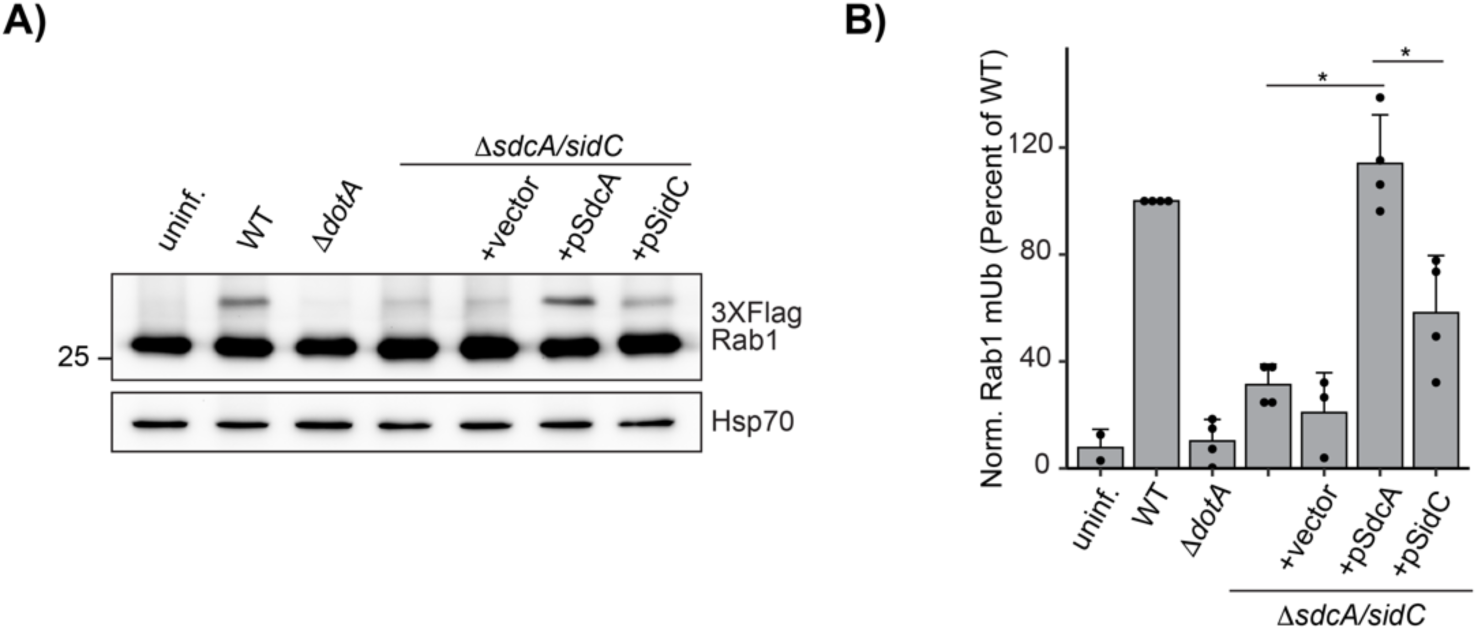
Rab1 ubiquitination is primarily promoted by SdcA. Related to Figure 7. (**A**) Immunoblot analysis of lysates prepared from HEK293T FcγR transfected with 3XFlag Rab1A and infected with a Δ*sidC/sdcA L.p*. strain panel (WT, Δ*dotA*, Δ*sidC/sdcA*, and Δ*sidC/sdcA* complemented with empty vector or plasmid encoded SdcA or SidC) for 1 hour (MOI=50). **(B**) Quantification of biological replicates (N=3) of experiment shown in (A). Normalized Rab1A monoubiquitination intensity was calculated as a percentage of WT *L.p.* infection levels (see Methods).

**Figure 7-S2:**
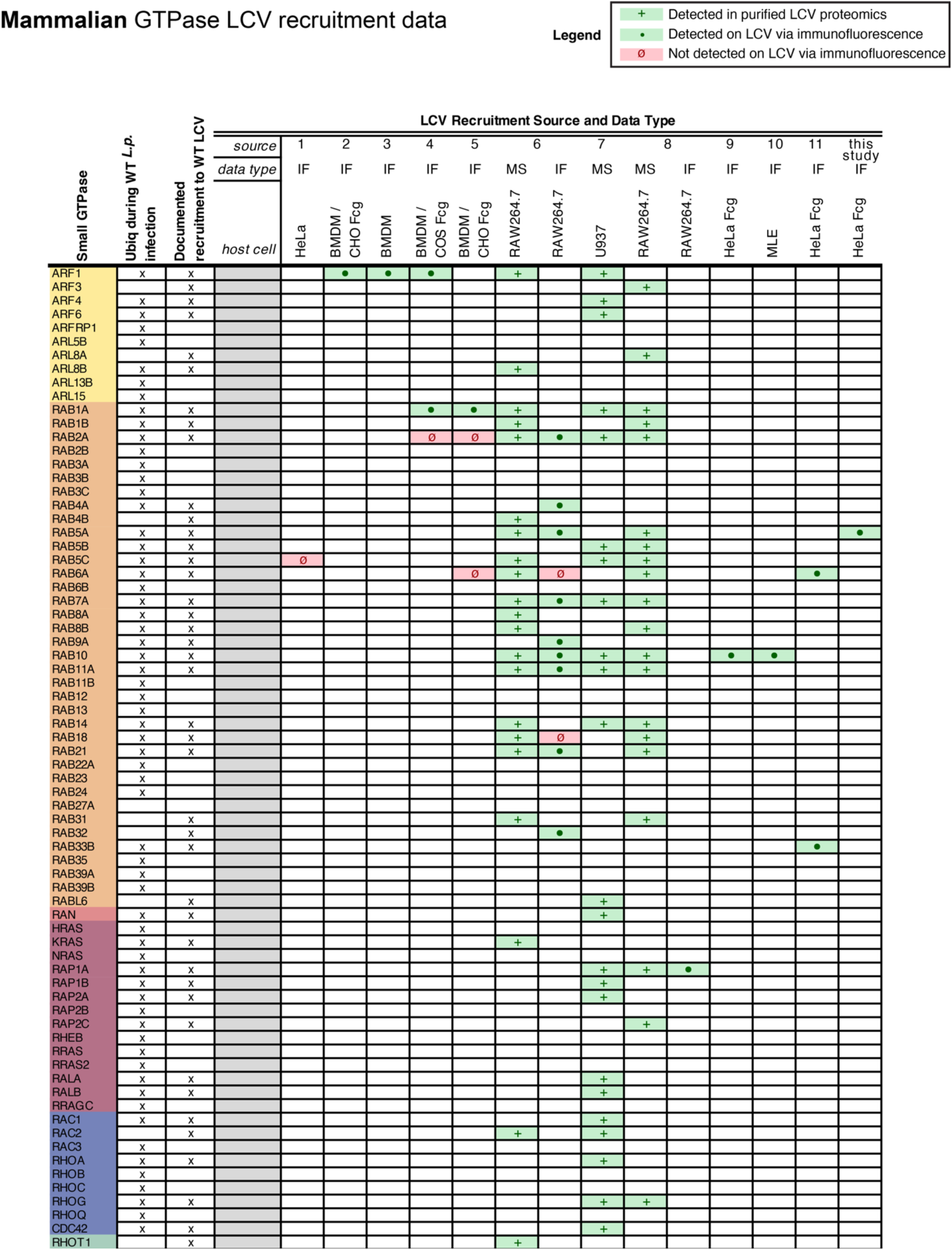
Small GTPases with detected ubiquitinations overlap with mammalian small GTPases known to be recruited to the LCV. Related to Figure 7. Compiled list of (1) small GTPases with detected ubiquitination in our ubiquitinomics data and (2) mammalian small GTPases known to be recruited to the LCV as assessed by immunofluorescent and/or purified LCV mass spectrometry approaches. Data type indicators: (+) detected via mass spectrometry of purified LCV, (•) detected via immunofluorescence, (ø) not detected via immunofluorescence. Sources: **(1**) (Clemens *et al*., 2000a) **(2**) (Kagan and Roy, 2002) **(3**) (Nagai *et al*., 2002) **(4**) (Derré and Isberg, 2004) **(5**) (Kagan *et al*., 2004) **(6**) (Hoffmann *et al*., 2014) **(7**) (Bruckert and Kwaik, 2015) **(8**) (Schmölders *et al*., 2017) **(9**) (Jeng *et al*., 2019) **(10**) (Liu *et al*., 2020) **(11**) (Kawabata *et al*., 2021).

**Figure 7-S3:**
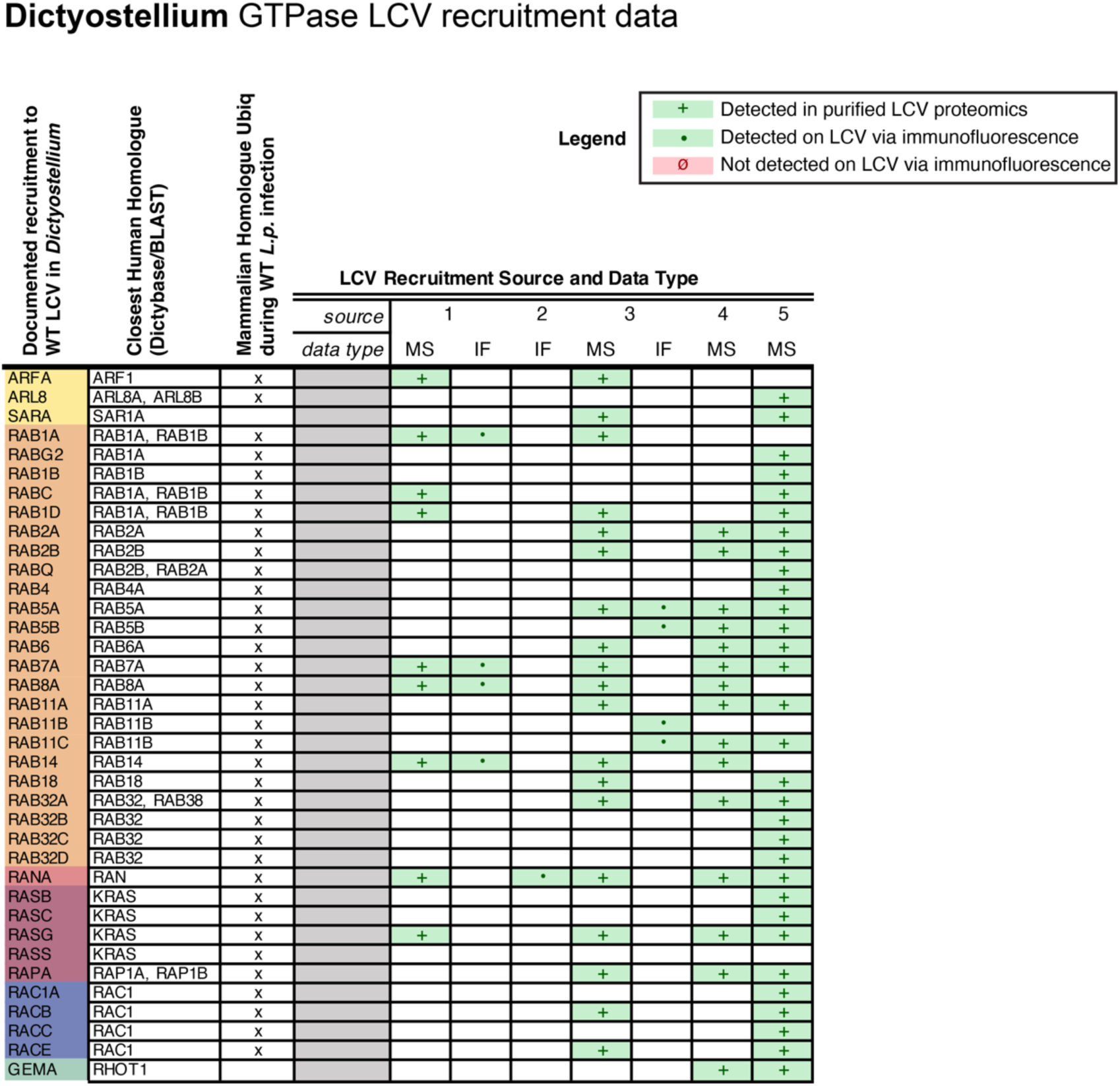
Small GTPases with detected ubiquitinations overlap with *Dictyostelium* small GTPases known to be recruited to the LCV. Related to Figure 7. Compiled list of *Dictyostelium* (amoebal) small GTPases known to be recruited to the LCV as assessed by immunofluorescent and/or purified LCV mass spectrometry approaches. Data type indicators: (+) detected via mass spectrometry of purified LCV, (•) detected via immunofluorescence, (ø) not detected via immunofluorescence. Closest human homologues were determined via Dictybase (http://dictybase.org/) and/or protein BLAST. Sources: **(1**) (Urwyler *et al*., 2009) **(2**) (Rothmeier *et al*., 2013) **(3**) (Hoffmann *et al*., 2014) **(4**) (Schmölders *et al*., 2017) **(5**) (Vormittag *et al*., 2023).

**Supplemental Table 1 – HPLC and MS methods.xlsx**

**Supplemental Table 2 – Infection Proteomics Dataset.xlsx**

**Supplemental Table 3 – Pathway and Protein Complex Enrichment Analysis.xlsx**

## Author contributions

Conceptualization, S.M., V.B., A.S., D.J-M., D.L.S., and N.J.K.; Methodology, S.M., E.S., G.M.J., D.J-M., D.L.S., and N.J.K.; Data Curation, A.S., V.B., D.J-M., D.L.S.; Validation, A.S., V.B.; Formal Analysis, A.S., V.B., D.J-M.; Investigation, V.B., A.S., T.M.; Writing, V.B., A.S., S.M., D.L.S., and D.J-M.; Visualization, A.S., V.B., D.J.-M.; Funding Acquisition, S.M., D.L.S., and N.J.K.; Resources, S.M., D.L.S., and N.J.K.; Supervision, S.M., D.L.S., and N.J.K..

## Competing interests

The N.J.K. laboratory has received research support from Vir Biotechnology, F. Hoffmann-La Roche and Rezo Therapeutics. N.J.K. has previously held financially compensated consulting agreements with the Icahn School of Medicine at Mount Sinai, New York and Twist Bioscience Corp. He currently has financially compensated consulting agreements with Maze Therapeutics, and Interline Therapeutics, Rezo Therapeutics, and GEn1E Lifesciences, Inc.. He is on the Board of Directors of Rezo Therapeutics and is a shareholder of Tenaya Therapeutics, Maze Therapeutics, Rezo Therapeutics and Interline Therapeutics. D.L.S. has a consulting agreement with Maze Therapeutics. All other authors declare no competing interests.

## Supporting information

Supplemental Table 1 HPLC and MS methods

Supplemental Table 2 Infection Proteomics Dataset

Supplemental Table 3 Pathway and Protein Complex Enrichment Analysis

## Acknowledgements

We thank Dr. Julia Noack for her preparation of the schematic in Figure 1A and for her support with initial analysis of the mass spectrometry data. We thank Dr. Philipp Schlaermann for preparation of cell pellets for the proteomics experiment. We thank Dr. Advait Subramanian and Dr. Joe Henry Steinbach for critically reading the manuscript. V.B. acknowledges support from the Moritz-Heyman Discovery Fund. A.S. acknowledges support from the UCSF iMicro Program MPHD T32 training grant. S.M. acknowledges financial support from the National Institutes of Health (R01GM140440, R01GM144378), the Pew Charitable Trust (A129837), Bowes Biomedical Investigator award, and a gift fund from the Chan-Zuckerberg Biohub. N.J.K. acknowledges financial support from the National Institutes of Health (U19 AI135990).

